# Surface color and predictability determine contextual modulation of V1 firing and gamma oscillations

**DOI:** 10.1101/421040

**Authors:** Alina Peter, Cem Uran, Johanna Klon-Lipok, Rasmus Roese, Sylvia van Stijn, William Barnes, Jarrod R Dowdall, Wolf Singer, Pascal Fries, Martin Vinck

## Abstract

The integration of direct bottom-up inputs with contextual information is a canonical motif in neocortical circuits. In area V1, neurons may reduce their firing rates when the (classical) receptive field input can be predicted by the spatial context. We previously hypothesized that gamma-synchronization (30-80Hz) provides a complementary signal to rates, encoding whether stimuli are predicted from spatial context by preferentially synchronizing neuronal populations receiving predictable inputs. Here we investigated how rates and synchrony are modulated by predictive context. Large uniform surfaces, which have high spatial predictability, strongly suppressed firing yet induced prominent gamma-synchronization, but only when they were colored. Yet, chromatic mismatches between center and surround, breaking predictability, strongly reduced gamma-synchronization while increasing firing rates. Differences between colors, including strong gamma-responses to red, arose because of stimulus adaptation to a full-screen background, with a prominent difference in adaptation between M- and L-cone signaling pathways. Thus, synchrony signals whether RF inputs are predicted from spatial context and may encode relationships across space, while firing rates increase when stimuli are unpredicted from the context.

## Introduction

Neuronal responses to sensory inputs are strongly modulated by the spatio-temporal context in which they are embedded. For instance, the firing rates of V1 neurons to stimuli in their (classical) receptive field (RF) can be increased or decreased by stimuli presented in their surround. Surround modulation is mediated by lateral and feedback connections, through which a given V1 neuron can be informed about a larger region of space than covered by its classical receptive field (Angelucci et al., 2017; Gilbert, 1992; Lund et al., 2003). Numerous related functions and computations have been linked to surround modulation, such as normalization (Carandini and Heeger, 2011), contour integration (Liang et al., 2017), perceptual filling-in (Land, 1959; Wachtler et al., 2003; Zweig et al., 2015), figure-ground separation (Lamme, 1995), computation of a saliency map (Coen-Cagli et al., 2012; Li, 2002), as well as efficient and predictive coding operations (Rao and Ballard, 1999; Vinje and Gallant, 2000). Theories of efficient coding postulate that surround suppression of neuronal firing may contribute to remove image redundancies across space from neuronal representations (Barlow, 2001; Coen-Cagli et al., 2012, 2015; Rao and Ballard, 1999; Schwartz and Simoncelli, 2001; Simoncelli and Olshausen, 2001; Vinje and Gallant, 2000; Zhu and Rozell, 2013). Theories of predictive coding in turn interpret the resulting neuronal response as a prediction error signal (Friston, 2005; Rao and Ballard, 1999; Spratling, 2010).

Recent work has extended the frameworks of efficient and predictive coding beyond firing rate modulations to include neuronal synchronization (Bastos et al., 2012; Chalk et al., 2016; Jadi and Sejnowski, 2014; Vinck and Bosman, 2016). Neuronal synchronization plays a functional role for the encoding and transmission of information, as well as for synaptic plasticity, and may therefore play an important role in contextual integration processes (Abeles, 1982; Akam and Kullmann, 2010, 2014; Azouz and Gray, 2000; Ballard and Jehee, 2011; Ballard and Zhang, 2018; Bernander et al., 1994; Borgers and Kopell, 2008; Bressler et al., 1993; Buzsaki, 2006; Buzsaki and Wang, 2012; Cardin et al., 2009; Fries, 2005, 2009; Fries et al., 2007; Havenith et al., 2011; Kempter et al., 1998; Kopell et al., 2000; O’Keefe and Recce, 1993; Palmigiano et al., 2017; Salinas and Sejnowski, 2000, 2001; Sejnowski and Paulsen, 2006; Singer, 1999; Singer and Gray, 1995; Sohaletal., 2009; Varela et al., 2001; Vinck et al., 2010a; Wang, 2010). A distinguishing feature of V1 activity, induced by many stimulus conditions, is synchronization of neuronal activity in the gamma-frequency band (≈30-80 Hz) (Fries, 2009; Gieselmann and Thiele, 2008; Gray et al., 1989; Jia et al., 2013b; Ray and Maunsell, 2010; Vinck and Bosman, 2016). A link between contextual modulation processes and V1 gamma-band synchronization is suggested by the finding that the amplitude of V1 gamma oscillations increases with stimulus size (Chalk et al., 2010; Gieselmann and Thiele, 2008; Gray et al., 1990; Jia et al., 2011, 2013b; Perry et al., 2013; Ray and Maunsell, 2011). There are different views on the way in which gamma oscillations might relate to predictive and efficient coding operations (Arnal and Giraud, 2012; Bastos et al., 2012; Chalk et al., 2016; Jadi and Sejnowski, 2014; Korndorfer et al., 2017; Vinck and Bosman, 2016). Bastos et al. (2012) and Arnal and Giraud (2012) hypothesized that gamma-band synchronization subserves the encoding and transmission of prediction error signals in the feedforward direction, and that lower frequency bands carry feedback predictions from higher areas (Arnal and Giraud, 2012; Bastos et al., 2012, 2015). Consistent with this hypothesis, bottom-up and top-down Granger-causal influences are strongest in the gamma and alpha/beta (≈10-20 Hz) band, respectively (Bastos et al., 2015; Bosman et al., 2012; Bressler et al., 2006; Richter et al., 2018; van Kerkoerle et al., 2014). According to this hypothesis, a mismatch between center and surround stimuli should lead to an increase in both firing rates and gamma-band synchronization, conveying prediction error signals.

In contrast, Vinck and Bosman (2016) recently hypothesized that (1) the amplitude of gamma oscillations in a given column reflects the extent to which classical RF inputs are predictable from the surround, and (2) that gamma-band synchronization among columns with non-overlapping RFs reflects predictability among their visual inputs. This could in turn provide a mechanism for orchestrating interactions between distributed neuronal columns, and for integrating efficiently-encoded signals in higher visual areas (Vinck and Bosman, 2016) (see Discussion). According to this hypothesis, redundancy between center and surround stimuli should lead to a decrease in firing rates (reflecting efficient coding) yet an increase in gamma-band synchronization. To distinguish between these conflicting views, precise manipulations of center-surround predictability are required.

As a starting point to test the interdependence between center-surround relationships, synchronization and firing rates, we considered the case of uniform surfaces and varied their size, center-surround relationships and color. The latter is an important feature for object recognition and visual search, and plays a role in social interactions and foraging (Bichot et al., 2005; Corso et al., 2016; D’Zmura, 1991; Gerald et al., 2007; Santos et al., 2001; Waitt et al., 2006). Furthermore, uniform surfaces are of particular interest because they contain highly redundant information across a relatively large image region. The predictability hypothesis (Vinck and Bosman, 2016) suggests that large, uniform surfaces should reliably induce gamma-band synchronization but be accompanied by low neuronal discharge rates. It further suggests that a modulation of center-surround predictability due to a chromatic mismatch (e.g. red center and green surround stimulus) should strongly reduce gamma-band synchronization but increase firing discharge rates.

Both the local color (chromatic) and luminance (achromatic) information of a surface could be extracted by visual cortex from two sources, namely either (1) by neurons with receptive fields at the uniform region of the surface (local information), or (2) by neurons with receptive fields at the edge of the surface (edge-derived information). An example of local color representation are the responses of single-opponent, hue-selective neurons (e.g. LGN and V1 cells with L+/M-, M+/L-, or blue (S) and yellow (L and M) RF sub-regions) (Livingstone et al., 1984; Shapley and Hawken, 2011). An example of local luminance representation are the transient responses of V1 neurons to temporal luminance changes, e.g. to flashed black or white stimuli (Shapley and Hawken, 2011; Zurawel et al., 2014).Edge-derived representations likely rely on an inductive “fill-in” process, in which neurons with RFs at the surface’s edge signal the difference between the color and luminance of the surface and the background. This may then change the firing of neurons with RFs at the surface’s center (Land, 1959; Wachtler et al., 2003; Zweig et al., 2015). The relative contributions of local (center) versus edge-derived representations remain largely unknown and likely differ between chromatic and achromatic surfaces (Zurawel et al., 2014; Zweig et al., 2015). Voltage-sensitive dye recordings of V1 populations, comparing the responses at the surface’s center to the edge, have shown that representations of achromatic surfaces depend more strongly on a “fill-in” process than chromatic surfaces (Zweig et al., 2015). Because contextual interactions likely have a different nature for achromatic than chromatic stimuli, we asked whether there are differences in the contextual modulation of firing activity and gamma-band synchronization between these two classes of stimuli.

Thus, we investigated the contextual modulation of V1 firing activity and gamma-band synchronization using chromatic and achromatic surfaces of different sizes, and a center-surround mismatch paradigm. Additionally, we examined differences among color hues, adaptation over time, and the influence of the full-screen background on which surfaces were displayed.

## Results

We recorded multi-unit (MU) activity and local field potentials (LFP) from the primary visual cortex (area V1) in two macaque monkeys, while they performed a fixation task. These recordings were made using a 64 channel chronic microelectrode array in monkey H and a 32-channel semichronic microelectrode array in monkey A (see Methods). Classical receptive fields (RFs, referring to classical RFs unless otherwise mentioned) of the MU activity were estimated using moving bar stimuli (see Methods; monkey H: median RF eccentricity 6.2 deg, range 5.2-7.1 deg; monkey A: median eccentricity 5.4 deg, range 3.2-8.5 deg).

We first studied LFP and MU responses to the presentation of stationary surface stimuli, namely large uniform disks covering the cluster formed by all RFs (6 deg diameter, flashed on and then maintained on screen; Figures 1A-B; *Dataset 1,* see Methods). The stimuli did not overlap with the fixation spot. Note that the stimuli were much larger than the RFs of the multiunits, such that they covered a large portion of the multi-units’ respective surround regions.

**Figure 1:**
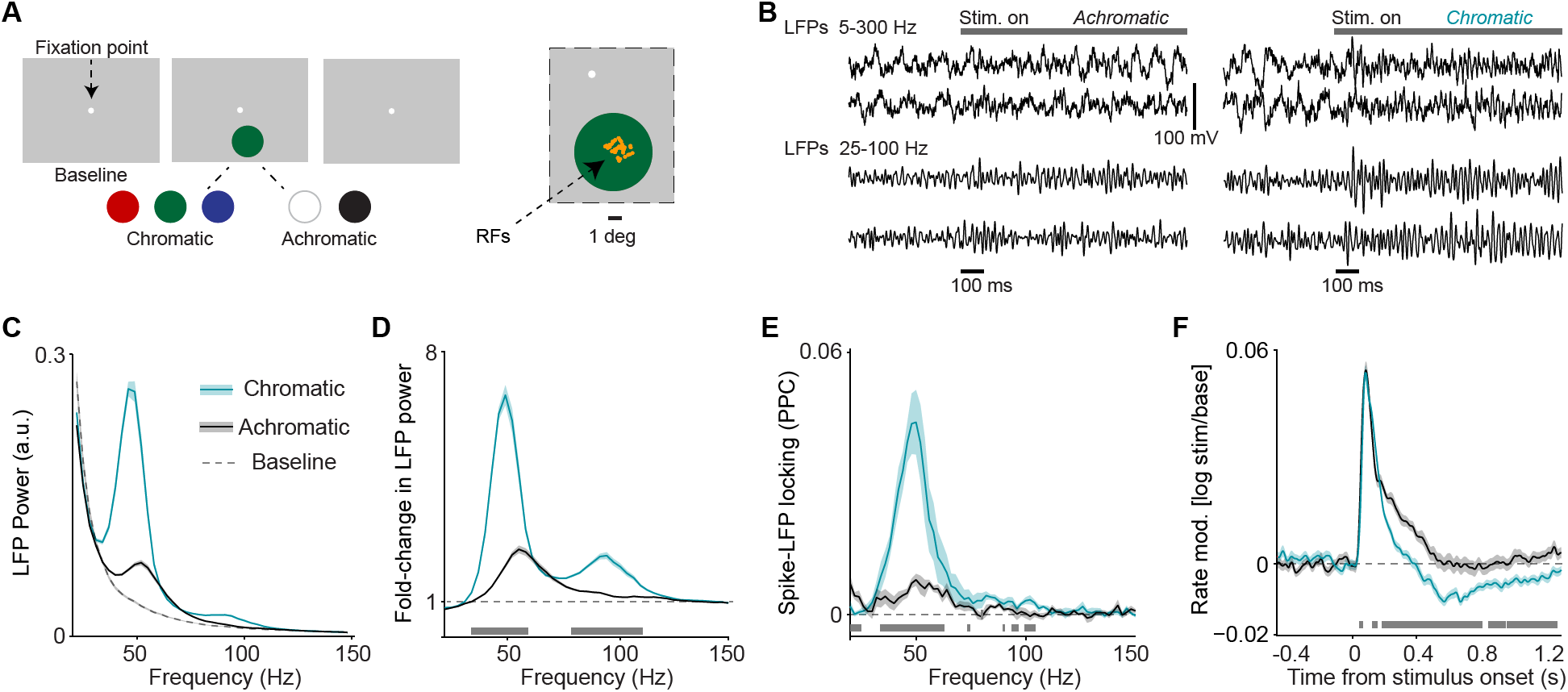
Analysis of LFP and multi-unit activity in response to large, uniform surfaces. (A) Illustration of experimental paradigm with large, 6 deg diameter surfaces (*Dataset* 1, see Methods; n = 5 sessions, 60±0 trials per session for chromatic and 40±0 trials per session for achromatic conditions. Trials numbers were euqated by random subselection for statistics.). Trials were self-initiated by fixating on the fixation spot (enlarged for visibility), followed by a baseline period of 0.5-0.8 s with a gray background screen. Surfaces were either chromatic (red, blue, or green) or achromatic (black or white) and presented for 3.3 s, the first 1.3 s of which are analyzed here. Right panel shows the RF locations of analyzed sites in one session. (B) Representative trials of LFP signals for achromatic and chromatic conditions (having gamma power close to the median of the respective condition). (C) Average LFP power spectra for chromatic (turqoise), achromatic (black) and baseline (gray) conditions. LFP power was estimated using Discrete Fourier Transform of non-overlapping epochs of 500 ms, with multi-tapering spectral estimation (±5 Hz). LFP spectra for all three conditions were normalized to the summed power (>20 Hz) for the baseline (gray) condition (see Methods). (D) Average change in LFP power, expressed as fold-change, relative to baseline. (E) Average MU-LFP locking, which was estimated using the pairwise phase consistency (PPC, see Methods). (F) Modulation of firing rate relative to baseline, expressed as *log*_*10*_*(stim/base).* (D-F) Shadings indicate standard errors of the means obtained with bootstrapping (see Methods). Gray bars at bottom of figure indicate significant differences between chromatic and achromatic stimuli, obtained from permutation testing with multiple comparison correction across all frequencies and time points (see Methods).

Initially, we analyzed differences between chromatic and achromatic surface stimuli (Figures 1 and 2; see Methods). We then considered the specific differences among responses to distinct color hues and achromatic stimuli (Figures 4-7). The LFP power spectra had similar frequency profiles in the two monkeys (i.e. the peaks were well aligned; Figure S2), and the MUs showed similar temporal profiles (Figure S2). Therefore, we pooled the data from the two animals. Note that statistical parameters are largely described in the figure captions.

**Figure 2:**
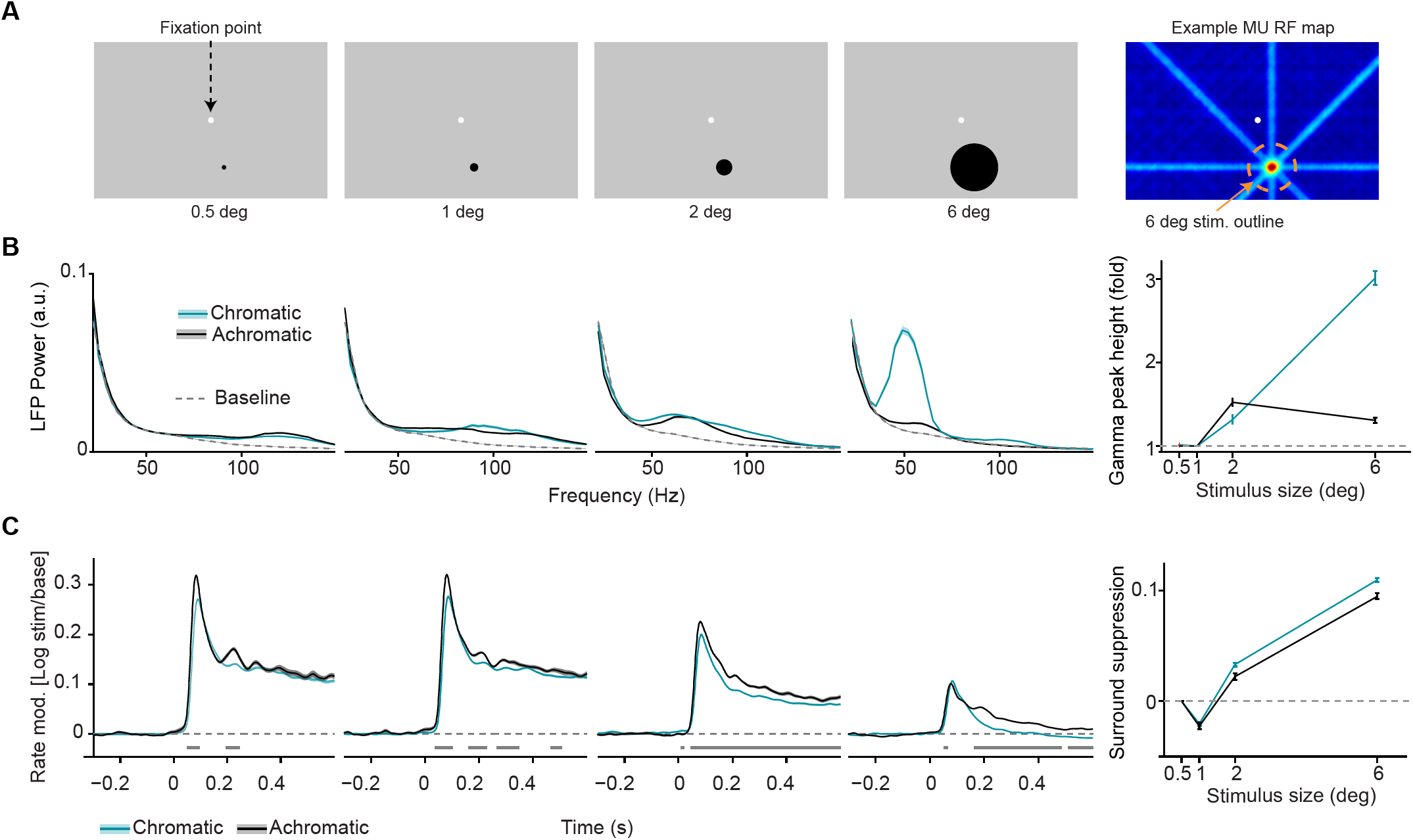
Dependence of LFP power spectra and MU firing activity on surface size. (A) Illustration of experimental paradigm (*Dataset* 2, see Methods; n = 9 sessions, 59.75±0.09/25.64±0.12 trials chromatic vs achromatic trials per condition in each session). Uniform surfaces of four different sizes were presented on a gray background screen. Fixation spot is enlarged for visibility. Right: Receptive field estimated with bar stimuli for a representative target channel, with the outline (orange dashed line) of the largest size stimulus (see Methods). Note that the activation outside the RF is due to the use of large bar stimuli sweeping over the monitor. (B) LFP power spectra for different sizes and chromatic/achromatic conditions. LFP power spectrum estimated and normalized as in Figure 1C, but now using 300 ms epochs. Right panel shows the gamma-band amplitude as a function of size, estimated using a polynomial fitting procedure between 30-80 Hz (see Methods). The difference between 6 and 0.5 deg stimuli was significantly larger for chromatic than achromatic condition (P<0.05, bootstrap test, see Methods). (C) Modulation of firing rate relative to baseline, expressed aslog_10_(*stim/base),* for different sizes and chromatic/achromatic conditions. Right panel shows surround suppression, which was defined as the difference in firing rate modulation between the 0.5 degree size and the other sizes.

### Characteristics of firing activity and LFP signals in response to uniform surface stimuli

We examined the effect of uniform surface stimuli on LFP power spectra. The presentation of large, chromatic surface stimuli (equiluminant red, green and blue; see Methods) induced prominent, narrow-band gamma oscillations in LFP power spectra (Figure 1C-D). These oscillations were clearly visible in the LFP traces (Figure 1B). In comparison, gamma-band oscillations were significantly weaker for achromatic surface stimuli (black or white, maximal contrast to background; Figures 1C-D). This finding was highly consistent across sites. Note that the differences between chromatic and achromatic stimuli do not result from differences in luminance contrast, which is addressed in detail in Figures 4 and S5. For each site and chromatic/achromatic condition, we determined the peak power change in the gamma-frequency range (30-80 Hz) using a polynomial fit (see Figure S1 and Methods). Gamma peak power changes were stronger for chromatic than achromatic surface stimuli at 97.8% (45 out of 46) of LFP recording sites (Figure S2).

To test whether V1 spiking activity was synchronized with the induced LFP gamma oscillations, we computed spike-field phase locking spectra (Pairwise Phase Consistency, Vinck et al. (2010b)) between MU and LFP activity obtained from nearby but separate sites (Figure 1E; see Methods). Spike-field phase-locking spectra for chromatic surface stimuli showed a prominent peak in the gamma-frequency band consistent with the gamma peak in the LFP power spectrum (Figure 1D), whereas phase-locking was significantly weaker for achromatic surface stimuli (Figure 1E).

Next, using the same stimulus paradigm, we examined the way in which the presentation of uniform surface stimuli affected MU firing activity. The presentation of chromatic and achromatic surface stimuli induced short-latency onset transients of similar magnitude (Figure 1F). However, we observed a stronger decrease in MU firing activity over time during continuous stimulus presentation for chromatic than achromatic surface stimuli, starting around 200 ms after the stimulus onset (Figure 1E). Strikingly, for chromatic surface stimuli, MU firing activity fell below baseline levels (Figure 1F). Note that in Figure 4D, we show that the decrease in MU firing below baseline only occurred for a subset of colors. The reduction in MU firing rates (0.3-1.3 s period) for chromatic as compared to achromatic surface stimuli was observed for 92% of recording sites.

These findings demonstrate that large, uniform, chromatic surface stimuli induce sparse yet highly gamma-synchronous V1 responses, whereas achromatic surface stimuli induce much weaker gamma-band synchronization but relatively more vigorous firing activity.

### Dependence of firing activity and LFP signals on stimulus size

The results shown in Figure 1 are consistent with the predictability hypothesis (Vinck and Bosman, 2016) outlined in the Introduction. Yet, they do not demonstrate directly that the enhancement in gamma-band synchronization observed for large uniform colored surfaces is due to contextual surround modulation, because we did not manipulate the surround input. Furthermore, it remains unclear whether the observed differences between chromatic and achromatic surfaces can be explained by a difference in contextual surround modulation or other factors like stimulus drive. To address these questions directly, we used a paradigm that varied the stimulus size across trials (Figure 2A; see Methods). We selected one site (or a few nearby sites with RF centers within 0.5 deg of the target site) per session and centered the stimulus on the multi-unit’s RF, which was previously mapped with moving bars. In each trial a stimulus of a particular size (0.5, 1, 2 or 6 deg diameter) was presented for 600 ms (Figures 2A-B).

We first examined how the characteristics of LFP power spectra depended on stimulus size. Analysis of LFP power spectra revealed a strong dependence of gamma power on stimulus size for chromatic stimuli, and by comparison a much weaker dependence for achromatic stimuli (Figure 2B). To quantify this size dependence, we determined the gamma peak power between 30-80 Hz (as described for Figure 1). For chromatic stimuli, increases in stimulus size resulted in increases in induced gamma peak power as soon as the stimulus also covered the surround (i.e. from 2 deg onwards, Figure 2B). By contrast, for achromatic stimuli, a gamma peak in the 30-80 Hz band emerged from 2 deg stimulus size onwards and showed no further increase with stimulus size. Given the relatively broad increase in >100 Hz LFP power seen in Figure 2B, we also determined gamma peak power and peak frequency in a wider range (30-150 Hz). This analysis revealed LFP power peaks >100 Hz for the sizes below 2 deg, and again the strong size dependence for chromatic compared to achromatic stimuli (Figure S3 A).

**Figure 3:**
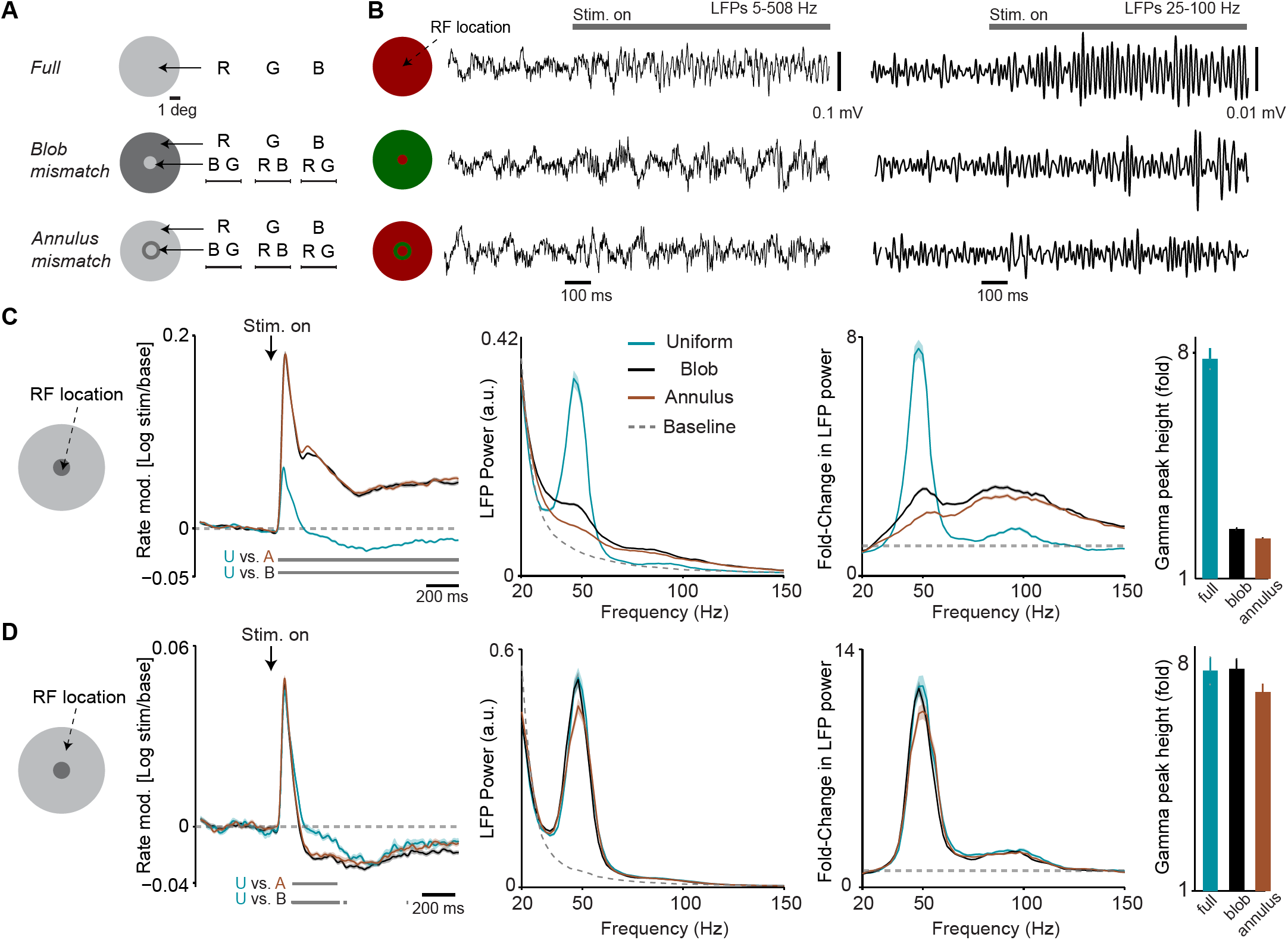
Dependence of LFP power spectra and firing rates on spatial predictability. (A) Illustration of paradigm (*Dataset 3,* see Methods; n = 8 sessions, 16.26±0.19 trials in each of the 15 conditions per session). We grouped stimuli intro three types. In the “Uniform surface” group of conditions, stimuli of 6 deg diameter were presented with either a red, blue or green hue (R B G, equiluminant). In the second “blob mismatch” group, the central 1 deg of the stimulus had a different (equiluminant) color than the rest of the stimulus. In the third “annulus mismatch” group, we presented an annulus ring (of 0.25 deg) of another color on top of the uniform surface (at equiluminant intensity) around the inner 1 degree from the stimulus center. All combinations of hues and stimulus types were presented, yielding a total of 15 individual conditions. (B) Representative LFP traces (having gamma power close to the median of all trials for the respective condition) for the three stimulus types. (C) Analysis for target channels with RFs at the center of the stimulus. Shown from left to right are: (1) The change in MU firing activity relative to baseline expressed as *log*_*10*_*(stim/base).* (2) LFP power spectra for the three stimulus conditions and the baseline. LFP power spectrum estimated and normalized as in Figure 1C. (3) The change in LFP power relative to baseline, expressed as a fold-change. (4) The gamma-band amplitude, estimated using a polynomial fitting procedure (see Methods). Gamma-band amplitude was significantly higher for uniform surface than blob and annulus conditions (P<0.05, bootstrap test, see Methods). (D) Same as (C), but now for target channels with RFs between 1.5 and 2 deg from the stimulus center, i.e. close to the central region of the larger, uniform region of the stimulus. Gamma-band amplitude did not significantly differ between conditions (bootstrap test, all P>0.08).

We further investigated the way in which MU firing was modulated by surround stimulation. We observed that for both achromatic and chromatic stimuli, MU firing activity was highest for 0.5-1 deg stimulus sizes (Figure 2C). This was consistent with the estimates obtained from RF mapping and the fact that we centered the presented stimuli on the MUs’ estimated RFs. For small stimuli (0.5-1 deg), only the initial transient in MU firing activity showed a difference between chromatic and achromatic conditions, with slightly higher firing activity for achromatic than chromatic stimuli (Figure 2C). In contrast, the presentation of a 2 or 6 deg stimulus, increasingly covering the surround, induced strong suppression of MU firing activity as compared to the 0.5 deg stimulus (Figure 2C, rightmost panel). This surround suppression was stronger for chromatic than achromatic stimuli (Figure 2C).

Furthermore, we analyzed responses during a later period in the trial, when the small stimulus had been presented for 600 ms, and a large (6 deg) surface stimulus of the same color was added for another 600 ms period (Figure S3). We found that this addition of the surround stimulus alone induced a rapid suppression of MU firing activity, which was significantly more pronounced for chromatic than achromatic stimuli (Figure S3).

These findings suggest that the relatively strong decrease in firing over time observed for large, chromatic surfaces (Figure 1) is at least partially explained by surround suppression. They furthermore indicate that for the small RF stimuli, there are no substantial differences on average between chromatic and achromatic surfaces in terms of MU and LFP responses. Yet, we find a prominent difference in the way chromatic and achromatic stimuli are affected by surround stimulation.

### Modulation of firing activity and LFP signals by center-surround predictability

A potential explanation for the results shown in Figure 1-2 may be the center-surround predictability hypothesis outlined in the Introduction (Vinck and Bosman, 2016). Yet, the employed paradigm used stimuli of different sizes, which may have recruited different neuronal circuits and may also have changed stimulus salience. We therefore used an additional stimulus paradigm in which surround influences were modified, while stimulus size was held constant. Specifically, we created three sets of equally sized stimuli. In one set, the surround was fully predictive of the RF stimulation, because it used a uniform surface (called “uniform” stimulus). In the second set (called “blob mismatch”), the surround was not predictive of the RF stimulation, because the surround stimulus and the 1 deg RF stimulus had different colors (which were physically equiluminant). In the third set (called “annulus mismatch”), the surround had the same color as the RF stimulation, but the two were separated by an annulus ring of a different, physically equiluminant color. This annulus ring had 0.25 deg thickness and an inner diameter of 1 deg.

We found that compared to the uniform surfaces, stimuli with a chromatic (blob or annulus) mismatch had higher MU firing activity (Figure 3C). This held true both for the initial transient period and the subsequent sustained response period (Figure 3C). At the same time, we observed a marked decrease in the amplitude of LFP gamma oscillations for the chromatic mismatch compared to the uniform surface stimuli (Figures 3B-C).

We further investigated whether this pattern of changes was specific to the sites having RFs near the center stimulus. To this end, we examined sites with RFs on the outer uniform regions of the stimulus (with RF centers 1.5-2 deg from the stimulus center; Figure 3D). For these sites the MU firing responses did not differ significantly between conditions during the initial transient period (Figure 3D). During the later sustained response phase, however, MU firing activity was reduced for the chromatic mismatch stimuli compared to the uniform surface stimulus (Figure 3D). Note that whenever the RF center covered a large uniform surface region, either in the uniform stimulus condition or when it covered the surround region of the mismatch stimuli, sustained firing levels were below baseline. This confirmed the respective finding reported in Figure 1.

These results suggest that a mismatch between stimuli at the RF center and the surround can dramatically change the surround influence on responses to the center. We wondered whether the surround influence on gamma oscillations originates from the uniform surface region or rather from the edge of the surface. To this end, we analyzed sessions in which we compared two sets of trials: First, trials with a full surface stimulus centered on a site’s RF (“RF-on-center” condition; Figure S4). Second, trials with a full surface stimulus positioned such that its edge fell into the RF center, i.e. with the surface shifted by 3 deg horizontally (“RF-on-edge” condition; Figure S4). We found that the amplitude of gamma oscillations was significantly higher at the center (“RF-on-center”) than at the edge of the surface stimulus (“RF-on-edge”), whereas the opposite was observed for MU firing activity (Figure S4). In one session (monkey H), we also showed disk stimuli that had their edge blurred with a Gaussian (2.5 deg size, 1 deg standard deviation). There were clear gamma responses also in this case (Figure S4).

Together, these results indicate that for colored surfaces the amplitude of gamma-band oscillations is commensurate with the “chromatic” predictability among visual inputs in space, and that gamma-band oscillations are not a mere consequence of input drive to a larger cortical region. Furthermore, these results suggest that gamma strength can be dissociated from stimulus salience, because the chromatic mismatch condition provided a highly salient stimulus in the RF, but resulted in weaker gamma.

### Differences in firing activity and LFP signals between color hues

The results above show prominent differences between chromatic and achromatic surfaces in terms of gamma-band synchronization. The respective analyses pooled different chromatic conditions (equiluminant red, green and blue) together. However, there may exist further differences in gamma-band synchronization within the chromatic conditions, i.e. between different hues. To investigate this we used two types of stimulus sets, which were presented in separate sessions. In the first stimulus set (Figure 4A) we presented each surface color at its maximum possible luminance level (given the limits of the employed monitor), and sampled from the entire spectrum of hues available with the monitor (see Methods). In the second stimulus set, we presented surface stimuli with different color hues at equated luminance levels (Figures 4B-C).

**Figure 4:**
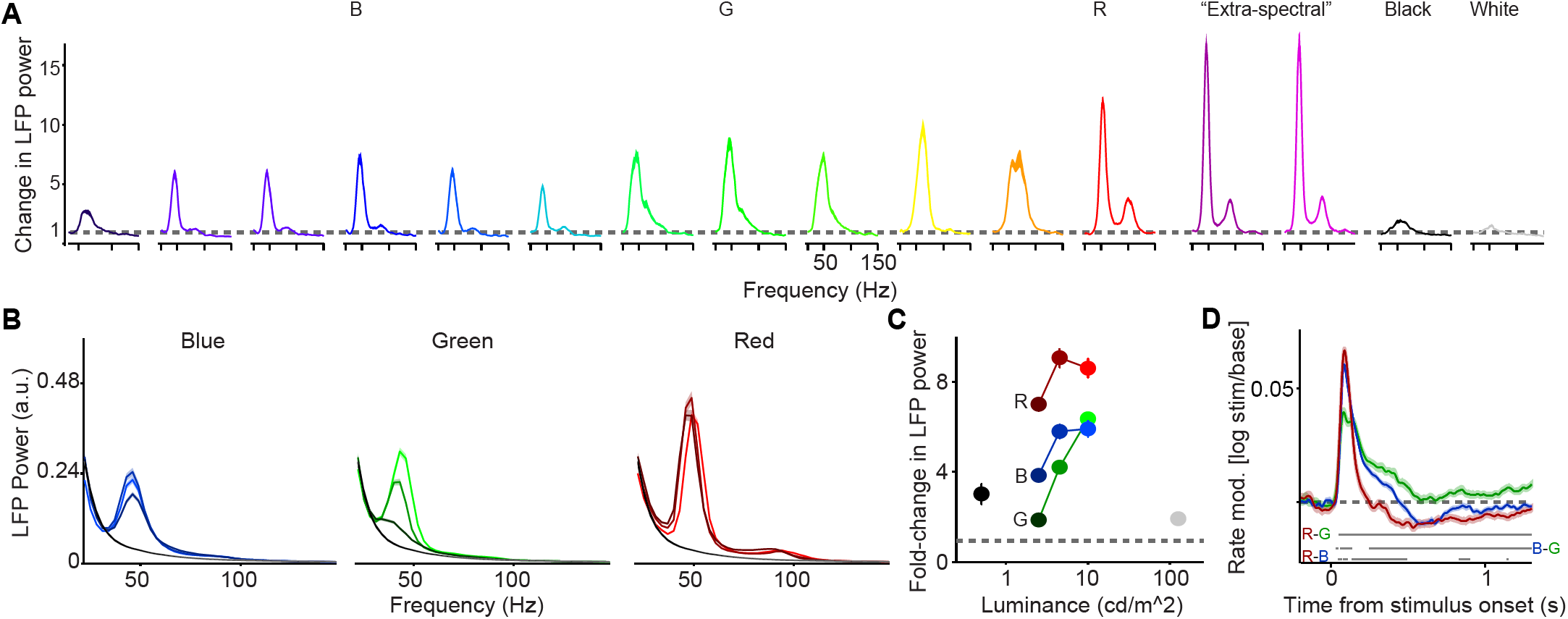
Dependence of LFP power spectra and MU firing activity on surface hue and luminance. (A) We presented uniform surfaces of 6 deg at maximum possible luminance levels, sampling from the available spectrum of wavelengths (*Dataset 4,* see Methods; n = 5 sessions, 15.24±0.1 trials per condition in each session). In addition, we presented black and white surfaces. Shown is the change in LFP power relative to the baseline (gray screen), expressed as a fold-change. (B) Three hues (red, green and blue) were presented at three different luminance levels (approximately 2.5, 5 and 10 cd/m^2^, *Dataset 1,* see Methods; n = 5 sessions, 19.8±0.45 trials per condition in each session). Shown are LFP power spectra. LFP power spectrum estimated and normalized as in Figure 1C. (C) Average change in LFP power, expressed as a fold-change, relative to the baseline (gray screen). The dependence of gamma amplitude on stimulus luminance was greater for G than for R or B (difference between high versus low: P<0.05, bootstrap test). Gamma oscillations amplitude R>B or G across all three luminance conditions, and B>G surface stimuli for low and intermediate luminance conditions (P<0.05, bootstrap test). (D) Modulation of firing rate relative to baseline, expressed as log_10_(stim/base). Horizontal bars at bottom of panel represent significant differences between stimuli at P<0.05 (permutation test, multiple comparison corrected for time bins).

Using the first stimulus set, we found that gamma-band LFP oscillations were reliably induced across the entire spectrum of hues (Figure 4A, Figure S5, see also Supplementary Table 1 for all luminance and CIE values). In addition, we found that gamma-band synchronization was reliably induced by surfaces with “extra-spectral” colors, i.e. colors resulting from a mixture of blue and red primaries (Figure 4A), as well as brownish hues. We further replicated our finding that gamma oscillations were relatively weak for both black and white surface stimuli as compared to all colored surfaces (Figure 4A).

For the first stimulus set (Figure 4A), the different colors were presented at their maximum possible luminance levels, which might confound the effects of hue and luminance. We therefore used a second stimulus set in which we presented surface stimuli with different color hues at three levels of equal physical luminance, i.e. different color values (Figures 4B-C). For all three hues, gamma amplitudes were greater for the highest compared to the lowest luminance condition (P<0.05, bootstrap test, see Methods; Figures 4B-C). The dependence of gamma amplitude on stimulus luminance was greater for green than for red or blue surface stimuli (difference between high versus low, Figures 4B-C). Gamma oscillations had a higher amplitude for red than for blue or green surface stimuli across all three luminance conditions (Figures 4B-C), whereas gamma amplitude was higher for blue than green surface stimuli for low and intermediate luminance conditions (Figures 4(B-C)). Another difference between the hues was that the gamma peak had a significantly lower frequency for green compared to red or blue surface stimuli (P<0.05, bootstrap test; Figures 4B-C and Figure S5B).

We further asked whether luminance-contrast modulated gamma oscillations and whether the observed difference between chromatic and achromatic conditions might be explained by luminance or luminance-contrast. A linear regression of gamma peak height against absolute Michelson contrast showed no significant relationship (p=0.16, F-test, Figure S5C). Furthermore, note that both in *Dataset 1* (Figures 1 and 4B) and *Dataset 4* (Figure 4A), there was relatively weak gamma for both black and white stimuli. Yet, compared to all the colored stimuli, the white stimulus was brighter and the black stimulus induced a greater luminance change relative to the gray background screen (see Methods and Table S1). These findings suggest that the difference between achromatic and chromatic surface stimuli observed in Figures 1-2 was not caused by chromatic surfaces being brighter or inducing a greater luminance change (contrast).

Given the relationships between MU firing activity and LFP gamma-band oscillations shown in Figures 1-3, we asked how these differences in LFP gamma oscillations were related to changes in firing activity. During the initial transient, MU firing activity was higher for red and blue rather than for green surface stimuli, with slightly stronger responses for red than blue surface stimuli (Figure 4D). Yet, we found that the post-transient decrease in MU firing activity over time was particularly pronounced for red and particularly weak for green stimuli (Figure 4D). In agreement with the data shown in Figure 1, we observed that MU firing activity fell below baseline levels for red and blue surface stimuli (Figure 4D).

Together, these results indicate that surfaces of all color hues tend to induce gamma-band oscillations with a higher amplitude compared to achromatic surfaces, and that the amplitude of gamma oscillations is relatively high for red surfaces.

### Temporal evolution of gamma-band responses

The observed differences in gamma oscillations between the color hues (Figure 4) might reflect a static and context-independent property of visual cortex to respond differently to distinct hues. Yet, the continuous presentation of a uniform surface stimulus for the duration of an entire trial likely induces substantial adaptation at many levels of the nervous system. We thus wondered whether different hues might adapt at different rates. To address this, we examined the temporal evolution of LFP power spectra over a longer time period, i.e. up to 3 s after stimulus onset. Time-frequency representations showed that qualitative differences between hues and between luminance levels tended to be relatively stable over time (Figure 5A). However, we found that the amplitude of gamma-oscillations decreased more rapidly over time for green than for blue or red surface stimuli (Figures 5A-B). This suggests that there may be differences in the time course and strength of adaptation between color hues, specifically stronger adaptation for green surface stimuli.

**Figure 5:**
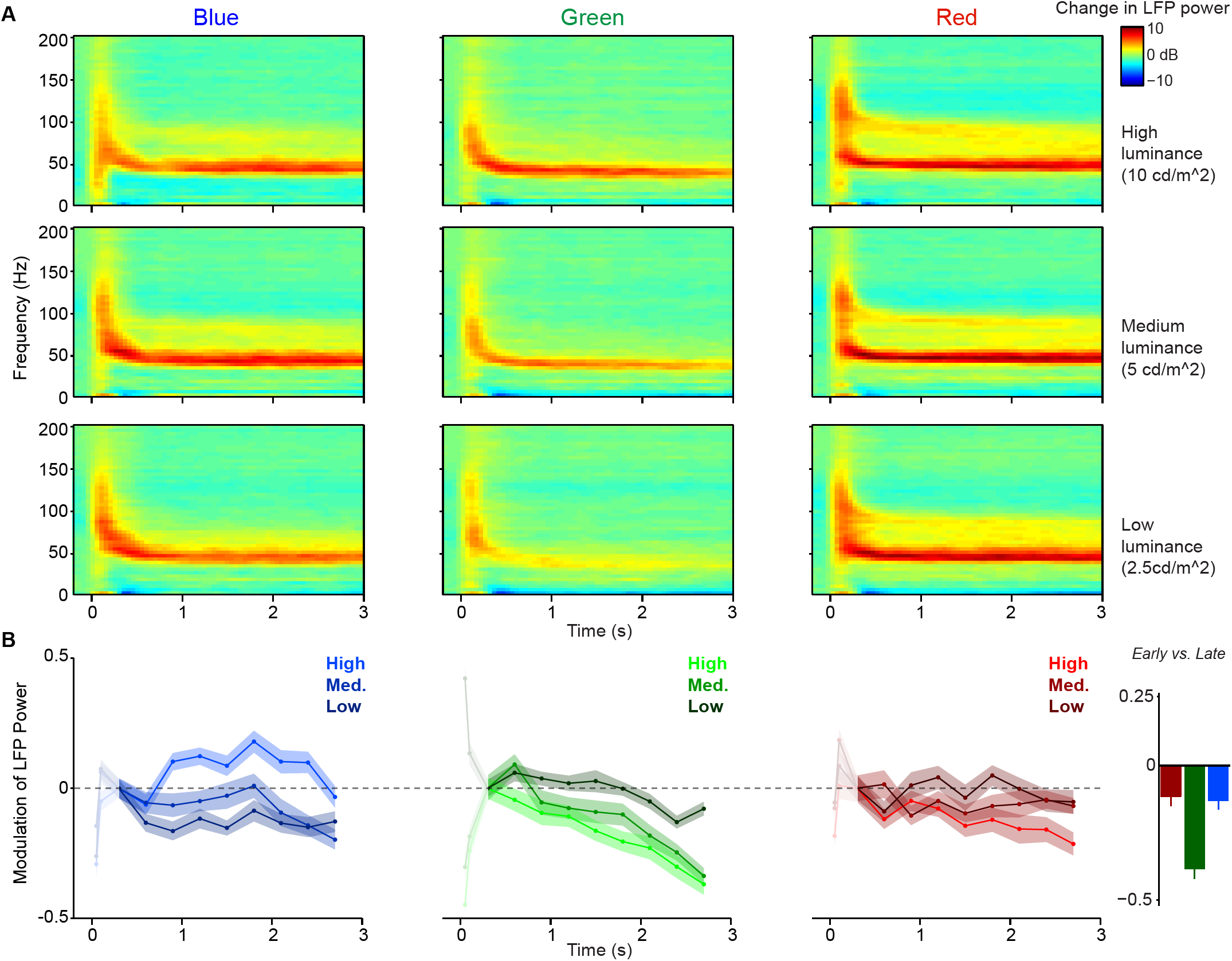
Within-trial temporal dynamics of LFP power spectra during viewing of uniform surfaces. Time-frequency representations of log-transformed change in LFP power relative to baseline (dB), using *Dataset 1* (n = 5 sessions, 19.8±0.45 trials per condition in each session). Shown are the three (equiluminant) different color hues at three different luminance levels. (B) Change in LFP power relative to 0.3-0.6 s period (third time point), separate for different color hues and luminance levels, shown as fold-change modulation index (see Methods). The first two points correspond to 0.05-0.35 and 0.1-0.4 s and are whitened out because these points are strongly influenced by initial firing transient and associated bleed-in of spiking activity at high frequencies. Right panel shows the modulation of LFP power in early (0.3-0.6 s) versus late (2.7-3.0 s) period, averaged over three luminance levels. The decrease in gamma peak amplitude over time was significantly larger for green than blue and red surfaces (peak amplitude estimated as described in Methods, main effect across luminance levels: P<0.05, bootstrap test), and did not differ between blue and red conditions. This also held true for the modulation of LFP power for the highest luminance condition only (P<0.05, bootstrap test).

### Dependence on full-screen background hue

One potential source of adaptation, other than the surface stimulus of a given trial, is the color composition of the continuously presented background. In the experiments described above, all surface stimuli were displayed on a gray full-screen background (FSB). Gamma-band responses to achromatic and chromatic surface stimuli may have been affected by the use of this gray FSB, given that the FSB itself may induce adaptation at many levels of the nervous system. We therefore asked how gamma-band responses to surface stimuli depend on the color of the FSB. To answer this question, we performed experiments in which we used different FSBs in separate, adjacent sessions (gray, white, black, blue, green, yellow and red) (Figure 6A; see Methods). The FSB was continuously presented during the entire session, i.e. remained on both during the pre-stimulus period, post-stimulus period and the period during which the surface stimuli were displayed (Figure 6A). In Figure 6, we analyze LFP responses to the presentation of chromatic surface stimuli of different hues, which were presented at the maximum possible luminance level (see Figure S6 for equiluminant red, green and blue as well as achromatic surface stimuli).

**Figure 6:**
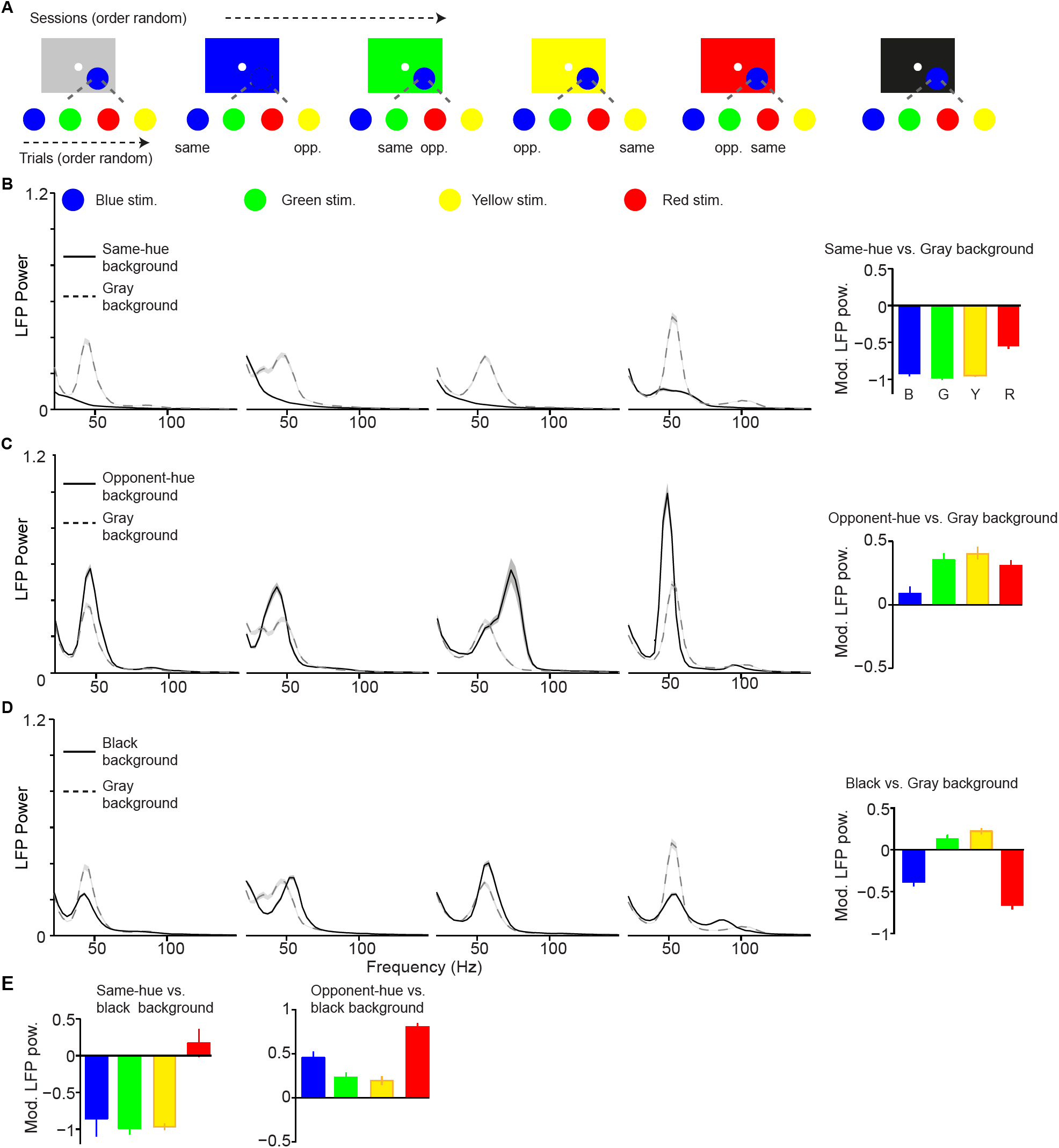
Dependence of LFP power spectra on background stimulus. (A) Illustration of paradigm (*Dataset 5,* see Methods; n = 25 sessions, 18.64±0.11 trials per condition and session). In a given session, a fixed background stimulus was used, and a set of chromatic and achromatic surfaces (6 or 8 deg) were presented in separate trials (see Methods). (B) Average LFP power spectra for the different color conditions during gray background versus same-hue background sessions. We show analyses for blue, green, yellow, and red surfaces, presented at maximum possible luminance. Right: modulation index of LFP gamma-amplitudes (see Methods). Main effect: P<0.05, bootstrap test. R versus G, B or Y, and G versus B: P<0.05, bootstrap test. (C) As B) for comparison of opponent color background and black background condition Main effect: P<0.05. B versus R, G or Y, P<0.05. (D) black background versus gray background. Main effect not significant. All color differences significant (P<0.05, bootstrap test). (E) Modulation of gamma-band amplitude for same-hue vs black background condition (left), as well as opponent-hue vs black background condition (right). Left: P<0.05: R versus G, B or Y; B versus G or Y Right: P<0.05: all combinations except G versus Y.

We first examined how the responses to surface stimuli with specific hues (e.g. green) were altered by using an FSB with the same hue (e.g. green), comparing them to the sessions with a gray FSB (Figure 6B). Surprisingly, we found that when the FSB had the same hue as the surface stimulus, there was a nearly complete abolishment of gamma oscillations for blue, green and yellow stimuli (Figures 6B and S6). This was observed both when the surface stimulus had the same luminance as the background (Figure 6) and when the surface stimulus had a lower luminance (Figure S6). Interestingly, red stimuli could still induce detectable gamma oscillations when presented on a red FSB, although the gamma amplitude was strongly reduced compared to the gray FSB condition (Figures 6B and S6).

The reduction in gamma-band oscillations for the same-hue FSB condition may have been an effect of stimulus size, because the background is effectively a very large surface. Alternatively, it may have been an effect of stimulus history. To investigate these possibilities, we analyzed the post-stimulus period immediately following the offset of a gray surface stimulus that was displayed on a colored FSB. We found that the reappearance of the FSB after the offset of the colored surface induced prominent gamma-band oscillations (Figure S6A). This indicates that the decrease in gamma-band oscillations with the same hue FSB condition was not due to the large size of the background color stimulation, but that it was due to the continuous presence of the same-hue background. This is also consistent with a previous report showing strong gamma with fullscreen color stimuli that change color across trials (Shirhatti and Ray, 2018), and with the positive relation between gamma and stimulus size shown in Figure 2C.

Next, we considered interactions between distinct hues. We wondered whether gamma oscillations can not only be reduced by same-hue FSBs, but also enhanced by FSB hues that are different from the stimulus hue, in particular when FSB and stimulus assume opponent colors. The organization of color vision around color-opponency axes, namely the red-green and the blue-yellow axes, is a key principle found both at the neurophysiological and psychophysical level (Livingstone et al., 1984; Solomon and Lennie, 2007; Tailby et al., 2008a; Wachtler et al., 2003). These color opponencies are thought to result from the computation of differences among signals deriving from L and M cones (red-green), and S cones versus L and M cones (blue-yellow). We found that for all surface hues, gamma oscillations were amplified when stimuli and FSBs were of opponent color hues (Figure 6C). This suggests that gamma oscillations are dependent on opponency signals along the red-green and the blue-yellow axes (Figure 6C).

Given the strong dependence of gamma-band oscillations on the FSB, we asked whether the use of a gray FSB may have induced differences in gamma-band amplitude among distinct hues. To examine this possibility, we used a black FSB, which should induce minimal adaptation for all cones. Quite surprisingly, the difference among red, green and blue hues that we had observed with a gray FSB could not be replicated when we presented the stimuli on a black FSB (Figure 6D). Compared to the gray FSB condition, gamma-band amplitudes were significantly lower for red and blue surface stimuli and significantly higher for yellow and green surface stimuli (Figure 6D). As a consequence, for the black FSB condition, gamma-band power was no longer highest in response to red stimuli, but showed a different dependence on hue (Figure 6D; Figure S6B). Specifically, gamma-band power was higher for green and yellow than red and blue surface stimuli (Figure 6D). The resulting pattern could not be explained by luminance contrast differences, because contrast increased for all hues on the black compared to the gray FSB, whereas gamma increased for some hues and decreased for others (Figure 6D).

In Figure 6B-C, we compared stimulus responses in the same-hue and opponent-hue background conditions with stimulus responses in the gray background condition. However, because of the evidence that the gray FSB may not have affected all stimulus hues equally, we also directly compared the same-hue and opponent-hue FSB conditions with the black FSB condition. This analysis revealed a marked difference between red and the other hues (Figure 6E). First, when an FSB of the same hue as the stimulus was compared to a black FSB, gamma was almost abolished for blue, green and yellow, but not for red stimuli. Second, when an FSB of the opponent hue was compared to a black FSB, gamma was enhanced for all colors, but particularly strongly for red (Figure 6E). The full matrix of gamma responses for different FSB conditions in Figure S6 shows that for all non-red chromatic FSBs, gamma oscillations were strongly amplified for red surface stimuli.

These data demonstrate that gamma oscillations depend strongly on the FSB, in a way that follows the color-opponency axes. Furthermore, a commonly used “default” of the display, namely gray, introduces adaptation effects that are color-specific.

### A quantitive model relating hue dependence of gamma-band oscillations to adaptation

To explain how gamma-band responses to surface stimuli depend on the FSB, we constructed a quantitative model by estimating the degree to which each FSB differentially adapts the S-, M- and L-cone pathways. Note that this model is agnostic to the neuronal locus at which adaptation of the distinct cone pathways occurs, e.g. it might occur in the retina, LGN or visual cortex. We hypothesize that gamma-band oscillations for colored surface stimuli are mediated by the activation of single color-opponent cells in a large spatial region by the same color input. The combination of bottom-up drive at each point of the surface and strong surround modulation may then lead to gamma oscillations. The result that gamma oscillations are particularly strong in the opponent-hue background condition (Figure 6) further suggests that when this circuit is more strongly activated (leading to stronger input drive as well as stronger surround modulation), gamma oscillations increase.

Following this reasoning, we further hypothesized that the dependence of gamma-band oscillations on the FSB can be explained by adaptation of specific cone pathways (see Discussion for further argumentation). As an example, a green FSB should lead to stronger adaptation of the M-cone compared to the L-cone pathway. This should increase the degree to which single-opponent cells with L+/M-RF subregions are activated by red surface stimuli, which may in turn increase the amplitude of gamma-band oscillations.

To capture this intuition in a quantitative manner, we constructed a model in which we aimed to predict the difference in gamma-band amplitudes between red and green surface stimuli (for blue and yellow surface stimuli see further below). The variable to be predicted was the red-green gamma ratio, defined as *γ*_*ratio*_ — log10 (*γ*_red_/*γ*_*green*_ ^), where^ *γ*_*red*_ and *γ*_*green*_ are the respective gamma-amplitudes for red and green surface stimuli. This *γ*_*ratio*_ was computed separately for all the different FSBs. We estimated the degree to which each FSB adapts the M- and L-cones, using the known response curves of the three cones as a function of wavelength from macaque monkeys (Harosi, 1987) (see Methods, Figure 7A). We then measured the physical wavelength spectrum for each FSB as realized on our monitor. We multiplied the FSB spectra of the different color primaries with the response functions of each cone and summed over wavelengths. This yielded for each FSB stimulus two parameter values, *M*_*adapt*_ and *L*_*adapt*_. We then fitted a multiple regression model predicting *γ* _*ratio*_ from *M*_*adapt*_ and *L*_*adapt*_ plus a constant regression intercept (Figure 7B). For this model, we used response data for both green and red surfaces presented at maximum possible luminance, as well as equiluminant red and green surfaces, across the different FSBs.

**Figure 7:**
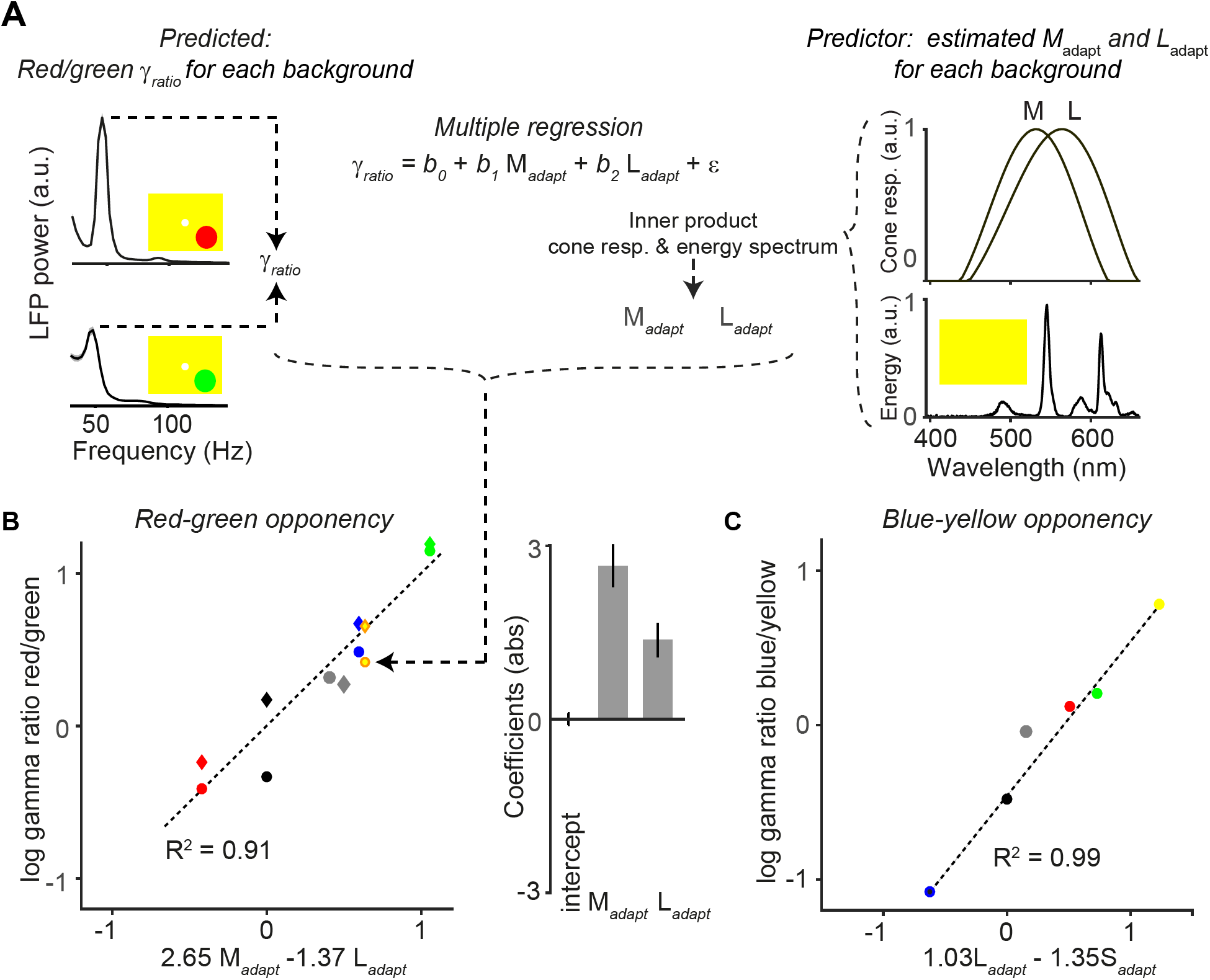
Quantitative model for dependence of gamma-band amplitude on background stimulus. (A) With a multiple regression model we predicted the ratio of gamma amplitude for red over green surface stimuli. Shown are the LFP gamma power spectra for red (top) and green (bottom) surface stimuli, with a yellow FSB. Normalized cone responses for macaque monkeys are shown on the right, constructed by fitting polynomials to bleaching difference spectra data (Harosi, 1987). For each full-screen background (FSB) we estimates the extent to which it adapted the three cones (*M*_*adapt*_, *L*_*adapt*_ and *S*_*adapt*_; see Methods), by convolving the spectral energy of the FSB with the normalized cone response curves. For illustration purposes we only show the M- and L-cone response curve. (B) The hues of the data points correspond to the FSB, and the shape indicates whether the red and green surfaces were presented at maximum possible luminance intensity (circle), or whether they were presented at equiluminant intensities (diamond). As predictor variables we used *M*_*adapt*_ and *L*_*adapt*_ for the different FSBs. The dependent variable was the red/green gamma-ratio, **γ**_*ratio*_. The coefficients indicates the relative influence of the two adaptation parameters on the **γ**_*ratio*_.,and the sign indicates whether adaptation of a given cone is increasing or decreasing the red/green gamma-ratio. (C) Similar to (B), but now for blue-yellow. In this case we used *S*_*adapt*_ and *L*_*adapt*_ as prediction parameters. A model using *M*_*adapt*_ in addition did not yield a significant coefficient for *M*_*adapt*_.

The regression analysis reveals that *γ*_*ratio*_ can be highly accurately predicted by the way in which each FSB adapts the L and the M cones (Figure 7B; R^2^ =0.91, *P <* 0.05, F-Test). The regression coefficients for *Madapt* and *Ladapt* were positive and negative, respectively. This indicates that adaptation of the M-cone increases *γ*_*ratio*_, whereas adaptation of the L-cone decreases *γ*_*ratio*_ (Figure 7B). We also found that the *γ*_*ratio*_ could not be significantly predicted when using *S adapt* and *L*_*adapt*_ as predictors (P=0.23), consistent with the idea that the neuronal mechanisms underlying the red-green opponency are dependent on the M versus L cone contrast. The regression intercept of the model (on the *γ*_*ratio*_ axis) was not significantly different from zero. This indicates that green and red tend to generate gamma oscillations of similar amplitude when the FSB does not adapt the cones, consistent with the findings shown for the black FSB (Figure 6). Strikingly, we found that the *M*_*adapt*_ coefficient had an absolute magnitude approximately twice as large as the *L*_*adapt*_ coefficient (Figure 7B). This suggests that uniform surfaces tend to adapt the M-cone pathway more strongly than the L-cone pathway, or that adaptation of the M-cone pathway has a stronger effect on gamma-band oscillations than adaptation of the L-cone pathway.

The model further explains some non-trivial findings that would have been unexpected if FSBs had affected the M- and L-cone pathway in a similar way: We observed that the yellow FSB strongly amplified *γ*_*ratio*_ (Figure 7B). Given its wavelength spectrum, the yellow FSB is expected to adapt the L-cones more strongly than the M-cones, which would predict a reduced *γ*_*ratio*_, i.e. red responses being weaker than green responses. By contrast, we found that *γ*_*ratio*_ was enhanced. The models explain this by the fact that the stronger L-cone than M-cone activation by the yellow background is more than compensated by the much greater *M*_*adapt*_ than *L*_*adapt*_ coefficient. Similarly, the *γ*_*ratio*_ increased for a gray compared to a black FSBs, even though gray FSBs should in principle adapt the M- and L-cone pathways to a similar degree (Figure 6). This was again compensated by the much greater *M*_*adap*_ than *L*_*adap*_ coefficient.

We performed a similar analysis for the yellow-blue (L+M - S) opponency axis, aiming to predict the gamma ratio of blue over yellow (Figure 7C). We first fitted a model with the S, L and the M cone parameters, and *γ*_*ratio*_ was now defined as *γ* _*ratio*_ = *log*_*10*_(*γ*_*blue*_*/*γ**_*yellow*_). This regression model explained a large degree of variance (R^2^ = 0.99), with almost equal magnitude of S (negative, -1.23) andL (positive, 1.24) coefficients, but a much smaller and non-significant coefficient for the M cone (-0.29). The finding that the model fit included a highly positive L-cone coefficient and non-significant (and negative) M-cone coefficient seems *prima facie* to contradict the canonical idea that the perceptual blue-yellow opponency axis is mediated by an S versus (L+M) opponency. However, neurophysiological data has shown that the main opponency for LGN cells on the yellow-blue axis is the L versus S cone (Tailby et al., 2008b). We further simplified our model using two predictive parameters, equating the blue axis to the S cone and the yellow axis to the L cone (Figure 7C). Again, we found a highly predictive relationship with negative weight for the S cone and a positive weight for the L cone (Figure 7C), with only a small and non-significant difference in the magnitude of the adaptation coefficients. These findings indicate that gamma oscillations are mediated not only by opponency signals along the red-green axis, but also along the blue-yellow axes.

## Discussion

### Summary

We investigated the way in which V1 responses to chromatic and achromatic surfaces are modulated by spatial and temporal context. We report the following main findings:

(1) Compared to achromatic surfaces, chromatic surfaces induced strong synchronization of neuronal activity in the gamma-frequency band (Figures 1 and 2). This finding held true for color hues across the entire wavelength spectrum (Figure 4).

(2) Whereas chromatic and achromatic surfaces induced an initial MU firing transient of similar magnitude (Figure 1 and 2), we found relatively weaker firing responses to chromatic surfaces in the sustained stimulation period, which was evident from a stronger decrease in firing over time related to an increase in surround suppression (Figures 1 and 2).

(3) Compared to uniform chromatic surfaces, composite stimuli with a chromatic mismatch between the stimulus covering the RF center and the surrounding surface evoked high firing activity, yet induced a very prominent reduction in the amplitude of gamma band oscillations (Figure 3). This supports the hypothesis of Vinck and Bosman (2016) outlined in the Introduction, namely that gamma-band synchronization reflects the degree to which inputs into the CRF can be predicted from the surround.

(4) Stimulus-induced gamma-band oscillations were also strongly modulated by the larger spatio-temporal context: We found that their amplitude depended strongly on the full-screen background (FSB) on which the surfaces were displayed (Figures 4-7). We concluded that the dependence of gamma-band synchronization on the FSB was explained by two key factors: First, color opponency along one of the two main color-opponency axes (red versus green and yellow versus blue). Second, a comparatively stronger adapting influence of FSBs on the M-cone pathway, which leads to comparatively stronger gamma oscillations for red surfaces for many (but not all) FSBs (Figures 6-7). This is consistent with our finding that the decrease in gamma-band amplitude within a trial is particularly strong for green surface stimuli (Figure 5).

### Differences between chromatic and achromatic surfaces

We asked whether there are differences in V1 gamma-band synchronization and firing responses between chromatic and achromatic stimuli. A previous voltage-sensitive dye imaging study has shown differences in V1 responses to chromatic compared to achromatic surfaces (Zweig et al., 2015). For achromatic surfaces, V1 response characteristics were indicative of a fill-in process, in which surface information emanates from the surface’s edge. This was not the case for chromatic surfaces (Zweig et al., 2015), which suggests that the V1 representation of surface color may largely depend on single-opponent LGN inputs to V1 and responses of V1 neurons with RFs within the uniform surface (Livingstone et al., 1984; Shapley and Hawken, 2011; Zweig et al., 2015). In the present study, we observed that chromatic surfaces exhibited much stronger gamma-band synchronization yet more suppressed firing than achromatic surfaces. This finding is consistent with the idea that V1 representations of surface color depend on the direct activation of neurons with RFs in the uniform region of the surface. It further supports the hypothesis that gamma-band synchronization arises from the predictability of visual inputs across space (Vinck and Bosman, 2016). Together, these data suggest that one could think of color, like stimulus orientation, as a “feature”, with predictive value for stimulus features in color space. If the features at the center stimulus are correctly predicted by the surround (context), the result is strong gamma-synchronization. On the other hand, predictability of stimulus luminance by itself in the absence of color information, as in the case of a large uniform white stimulus, may not be sufficient for inducing strong gamma-band synchronization, although stimulus luminance can further modulate gamma-band synchronization. In contrast, prominent gamma-band synchronization can be generated in response to achromatic stimuli when they have structural features (orientation, frequency, phase) that are highly predictable over space, e.g. bars and gratings (Chalk et al., 2010; Gieselmann and Thiele, 2008; Gray et al., 1989).

### Dependence of gamma-band oscillations on hue and full-screen background

We demonstrated a prominent difference in gamma-band synchronization and firing activity between chromatic and achromatic surfaces. However, we also found prominent hue-related differences. These differences could reflect a constant property of the visual system to respond differently to particular hues, or might arise from other contextual processes, such as adaptation to the full-screen background (FSB) on which the surfaces were displayed. When using a gray FSB, we found that gamma oscillations were particularly strong for surface stimuli with red hues. This finding is consistent with previous work that used a gray FSB throughout and showed stronger gamma-band synchronization for red stimuli (Rols et al., 2001; Shirhatti and Ray, 2018). Note that with a gray FSB, gamma oscillations were reliably induced by all surface hues, which is consistent with our findings suggesting that both yellow-blue and red-green opponencies contribute to the generation of V1 gamma oscillations (Figures 6 and 7).

Importantly, we found that differences in gamma-oscillation strength among hues were highly dependent on the FSB and that with a black background a different dependence on hue emerged. In particular, with a black compared to a gray FSB, gamma oscillations increased in amplitude for green and yellow surface stimuli, but decreased in amplitude for red and blue surface stimuli. The quantitative model presented in Figure 7 suggests that the M-cone pathway adapts more strongly than the L- and S-cone pathways, even when the adaptation-inducing FSB is supposedly “neutral” (gray). By extension, background hues during natural vision would play a similar adapting role. An explanation for this phenomenon may be that, in general, uniform surfaces induce stronger or faster adaptation of the M-cone than L- and S-cone pathways, which may have a retinal, thalamic and/or cortical source. This interpretation is consistent with our finding that gamma-band amplitude decreased on a time-scale of seconds more rapidly for green than red or blue surfaces (Figure 5). We observed several other unique response features for green surfaces consistent with the idea of differential M-cone adaptation: First, we found that gamma oscillations were strongly dependent on luminance for green surfaces in particular, which suggests that a stronger luminance is needed to overcome adaptation (Figure ??). Second, we found that gamma oscillations had a significantly lower peak frequency for green than for red or blue surface stimuli (Figure ??). This suggests a weaker stimulus drive for green than red surface stimuli, because enhancing stimulus drive has been shown to increase the frequency of gamma oscillations (Hadjipapas et al., 2015; Henrie and Shapley, 2005; Jia et al., 2013b; Ray and Maunsell, 2010; Roberts et al., 2013). Third, with a gray FSB, we found that evoked MU transients were weaker for green than for equiluminant red and blue surface stimuli, which is consistent with increased adaptation of the M-cone pathway. Yet, we found that green stimuli exhibited a weaker decrease in firing over time (Figure 4D), and that the decrease in firing over time was particularly pronounced for red stimuli.

These findings have two important implications: (1) A gray FSB may differentially change neuronal responses to surfaces of different color hues. This may have implications for the design of studies examining differences in neuronal or behavioral responses between color hues. (2) Differential adaptation to distinct hues may have consequences for color perception in general. Interestingly, at the psychophysical level, it has been shown that psychophysical after-effects emerge more rapidly after viewing green than viewing red stimuli (Werner et al., 2000). Differential adaptation of M-cones may reflect important behavioral requirements of primates. The visual environment of primates is dominated by green and yellowish stimuli like leaves and trees (Mizokami et al., 2003). Detection of fruits with relatively high energy in red hues may be an important behavioral task for many primates (Melin et al., 2017). Trichomacy provides behavioral advantages for such detection (Melin et al., 2017), which may be aided by fast adaptation of M-cone derived signals.

### Differential effects of spatial and temporal context on gamma-band synchronization

This study reports two main findings: When the RF stimulus is part of a larger uniform surface, the spatial context allows a prediction of RF content, and gamma oscillations are enhanced. Yet, when the RF stimulus is part of a longer uniform stimulation period, the temporal context allows a prediction of RF content, and gamma oscillations are reduced. This suggests that the two effects are brought about by different mechanisms. When RF content is spatially predictable, enhanced gamma oscillations are accompanied by reduced firing rates. This pattern of results is highly suggestive of enhanced inhibition (Vinck and Bosman, 2016), and is in line with the prominent role of inhibition in the generation of gamma (see *Mechanisms ofgamma-band synchronization* further below). By contrast, when the RF content is temporally predictable, the pattern of results is more suggestive of an adaptation mechanism leading to a progressive reduction in gamma strength. This part of our results is in line with previous reports from crossmodal and auditory studies (Arnal et al., 2011; Todorovic et al., 2011), which found enhanced gamma oscillations for unexpected stimuli. These findings are consistent with an earlier hypothesis, which stated that gamma oscillations should be enhanced for stimuli that generate prediction errors (Bastos et al., 2012). Note that the effects of both spatial and temporal predictability do not necessarily rely on top-down feedback. The relationship of V1 gamma-band synchronization to temporal context may be more complex than suggested by this general conceptual notion, however: Previous studies have shown that when continuous stimulus motion has a large degree of jitter/randomness, either in case of entire video frames or in case of bar stimuli, V1-gamma-band synchronization tends to be weak, whereas V1 gamma-band synchronization tends to be strong in cases where stimulus motion is predictable (Kayser et al., 2003; Kruse and Eckhorn, 1996; Vinck and Bosman, 2016). Thus, the notion that V1 gamma-band synchronization increases when stimuli are unexpected or salient given the temporal context might apply only to discrete stimulus onsets and not generalize to cases where there is continuous stimulus motion. Furthermore, stimulus repetition can lead to a monotonic increase in V1 gamma-band synchronization over trials (Brunet et al., 2014). This increase of V1 gamma-band synchronization with stimulus repetition may reflect a slower learning process in which spatial center-surround interactions are modified by experience (Vinck and Bosman, 2016).

### Mechanisms of gamma-band synchronization

The results discussed above revealed several principles underlying the stimulus dependence of gamma-synchronization. Yet, it remains unclear what precise neuronal mechanisms account for the emergence of V1 gamma-synchronization, and its dependence on center-surround predictability. The interaction between inhibitory and excitatory neurons likely plays a critical role (Bartos et al., 2007; Borgers and Kopell, 2005; Bush and Sejnowski, 1996; Buzsaki and Wang, 2012; Cardin et al., 2009; Csicsvari et al., 2003; Hasenstaub et al., 2005; Jadi and Sejnowski, 2014; Kopell et al., 2000; Lytton and Sejnowski, 1991; Moore et al., 2010; Perrenoud et al., 2016; Salkoff et al., 2015; Siegle et al., 2014; Sohal et al., 2009; Tiesinga and Sejnowski, 2009; Veit et al., 2017; Vinck et al., 2013a,b; Wang, 2010; Wang and Buzsaki, 1996; Whittington et al., 1995). Specialized electrophysiological sub-classes of pyramidal neurons like chattering (fast-rhythmic bursting) cells, which have resonant properties in the gamma-frequency band, could also be a critical component of gamma rhythmogenesis (Cardin et al., 2005; Gray and McCormick, 1996; Nowak et al., 2003). Tangential, excitatory connections linking preferentially columns with similar feature preferences (e.g. color or orientation) may play a crucial role in synchronizing neuronal assemblies coding for related features (Gray et al., 1989; Korndorfer et al., 2017; Vinck and Bosman, 2016). A stimulus with high spatial predictability is likely to simultaneously activate a large number of preferentially coupled columns. This could then give rise to enhanced cooperativity among these columns and boost gamma-synchronization by the recruitment of local excitatory and inhibitory neurons. In addition, feedback from higher visual areas could be critical, considering that the spatial spread of tangential connections is somewhat limited and covers a smaller surround region than cortical feedback (Angelucci et al., 2017).

### Functions of gamma-band synchronization

We finish with a discussion of the functional implications of the present findings. Early theories of gamma-synchronization proposed that it may contribute to solving the “binding problem” (Singer, 1999; Singer and Gray, 1995). This refers to the problem that the visual system segments images into segregated objects, which raises the problem that the local features comprising the object must at some processing stage be bound together. It was proposed that the activity of distributed neurons can be dynamically grouped together through synchrony according to perceptual Gestalt principles (Engel et al., 1992; Milner, 1974; Singer, 1999, 2018; Singer and Gray, 1995; Von Der Malsburg, 1994). Notably, functions that have been linked to surround modulation, such as contour integration (Liang et al., 2017), perceptual filling-in (Land, 1959; Wachtler et al., 2003; Zweig et al., 2015), and figure-ground segregation (Lamme, 1995), may contribute to perceptual grouping and underlie some of the Gestalt principles. Later work emphasized that gamma-synchronization can flexibly regulate communication between neuronal populations (Akam and Kullmann, 2010; Colgin et al., 2009; Fries, 2005; Jia et al., 2013a; Knoblich et al., 2010; Palmigiano et al., 2017; Salinas and Sejnowski, 2001). For example, the communication-through-coherence hypothesis states that communication between neuronal populations can be flexibly modulated by selective coherence according to cognitive demands (Fries, 2005, 2015). Recent studies have shown that neuronal groups in distant visual areas show gamma-band coherence primarily when they processes an attended stimulus and that the level of coherence predicts behavioral benefits of attention (Bosman et al., 2012; Buschman and Miller, 2007; Gregoriou et al., 2009; Grothe et al., 2012; Rohenkohl et al., 2018).

In the context of efficient and predictive coding and the relationship of V1 gamma with spatial predictability, V1 gamma-synchronization may play two functional roles (Vinck and Bosman, 2016), which remain to be tested:

1) Gamma-synchronization may be a mechanism to increase the effective synaptic gain of V1 neurons on post-synaptic targets (e.g. V2) (Bernander et al., 1991; Fries, 2005; Konig et al., 1996; Salinas and Sejnowski, 2000, 2001; Softky, 1994) when a stimulus is efficiently encoded. This may ensure reliable transmission of V1 outputs even when firing is sparse, which is especially important in the presence of noise within or competing inputs to the receiving area.

2) Gamma-synchronization could play an important role in coordinating the interactions between distributed V1 columns receiving related, and thereby redundant, visual inputs. The outputs of these columns need to be synaptically integrated, for which gamma-synchronization could be a mechanism (Fries, 2005; Konig et al., 1996)

In sum, the present work provides evidence that visual cortex shows sparse and gamma-synchronized responses when surround stimulation predicts RF center stimulation. In contrast, firing rates are high when the surround does not predict the center. These effects are particularly pronounced in case of chromatic, compared to achromatic surfaces. A second key insight is that the FSB on which surfaces are displayed strongly modulates gamma-synchronization, in a way that suggests that uniform surfaces lead to stronger adaptation of the M-cone compared to L-cone pathways. This not only explains differences in gamma-band oscillations between surfaces of different hues, but may also have important behavioral and perceptual consequences, which needs to be explored in future work.

## Acknowledgements

AP, CU and MV conceived of the idea of the study and designed the experiments. AP, CU, JKL and RR performed recordings. AP, JKL, RR, SS and WB performed initial behavioral training. JKL, SS, WB, and WS planned and performed surgical implants. For this we are also thankful to Michael Schmid and Richard Saunders. AP, JKL, JRD, WS and PF collected preliminary (unpublished) data on hue differences; in this context we would also like to acknowledge Gareth Bland and Marieke Scholvinck. AP, CU and MV performed data analysis. AP, CU, WS, PF and MV wrote the paper, with help from comments of the other authors. We would like to thank Quentin Perrenoud for very helpful comments. PF acknowledges grant support by DFG (SPP 1665, FOR 1847, FR2557/5-1-CORNET, FR2557/6-1-NeuroTMR), EU (HEALTH-F2-2008-200728-BrainSynch, FP7-604102-HBP, FP7-600730-Magnetrodes), a European Young Investigator Award, NIH (1U54MH091657-WU-Minn-Consortium-HCP), and LOEWE (NeFF). WS acknowledges the Reinhart Kosselleck grant of the German Research Foundation. We also wish to acknowledge Emmy Noether 2806 to Michael Schmid.

## Methods

All procedures complied with the German and European regulations for the protection of animals and were approved by the regional authority (Regierungsprasidium Darmstadt).

### Surgical procedures

Two male adult macaque monkeys (Macaca mulatta) were used in this study (age 9-10 years, 15-17 kg). All surgeries for implantations were performed under general anesthesia and were followed by analgesic treatment post-operatively. A head post was implanted in both monkeys to allow for head fixation. In monkey H, we implanted CerePort (”Utah”) arrays with 64 microelectrodes (inter-electrode distance 400 *μ*m, tip radius 3-5 *μ*m, impedances 70-800 kOhm at 1000 kHz, half of them with a length of 1 mm and half with a length of 0.6 mm, Blackrock Microsystems). One such array was implanted into area V1, another one in V4, both in the left hemisphere. The V4 array is not considered here. For array implantation, a large trepanation covering both areas was performed, the dura was cut open and reflected, arrays were inserted using a pneumatic device (Blackrock Microsystems), and both dura and bone were surgically closed. A reference wire was inserted under the dura towards parietal cortex. In monkey A, we implanted a semichronic microelectrode array Microdrive into area V1 of the left hemisphere (SC32-1, Gray Matter Research, containing 32 independently movable Alpha Omega glass insulated Tungsten electrodes with an impedance range of 0.5-2 MegaOhm and an inter-electrode distance of 1.5 mm). The microdrive chamber was used as the reference during recordings.

### Behavioral task

Both monkeys were trained on a fixation task. Monkeys were seated in a custom-made primate chair in a darkened booth. The two animals were positioned 83 (monkey H) or 64 cm (monkey A) in front of a 22 inch 120 Hz LCD monitor (Samsung 2233RZ, (Ghodrati et al., 2015; Wang, 2011). Both monkeys self-initiated trials by fixating on a small fixation spot, which was presented at the screen center. Monkey H performed a pure fixation task. For monkey H, the fixation spot was a Gaussian with a white center, tapering smoothly into the background. For recordings with white background, the fixation spot color was changed to red. Note that the pattern of results for gray and white FSBs was very similar despite this difference (Figure S6 B), and that receptive fields were not covering the fovea. The task of monkey A was to report a change in the fixation spot from red to green or blue (randomly) with a lever release. The change in the fixation spot occurred only after the stimulus period and an additional 700 ms of background stimulation, during which the animal maintained fixation. For the recordings with colored backgrounds in monkey A, fixation colors were changed to remain visible, with a magenta fixation spot during the baseline and stimulus period. For both animals, trials during which the eye position deviated from the fixation spot by more than 0.8-1.5 visual deg radius were aborted. Correctly performed trials were rewarded with diluted fruit juice delivered with a solenoid valve system.

### Recordings

Data acquisition was performed using Tucker Davis Technologies (TDT) systems. Data were filtered between 0.35 and 7500 Hz (3 dB filter cutoffs) and digitized at 24.4140625 kHz (TDT PZ2 preamplifier). Stimulus onsets were recorded with a custom-made photodiode. Eye movements and pupil size were recorded at 1000 Hz using an Eyelink 1000 system (Eyelink Inc.) with infrared illumination. Eye signals were calibrated before each recording session using a standardized fixation task.

### Visual stimulation paradigms during recordings

For all paradigms, stimuli were circular, did not have overlap with the fixation spot, and typically spanned a region from ca. 3-9 deg of eccentricity (monkey H) or 2.5-8.5 deg (monkey A, maximum: 1.6-9.6 deg for *Dataset 4*) in the lower right visual quadrant, matching RF locations. Trials always started with a baseline that lasted 0.5-0.6 s (monkey H) or 0.5-0.8 s (monkey A), and during which only the FSB and the fixation spot was shown. We used the following stimulus paradigms:

#### Dataset 1

For Figure 1, 4B and 5, we presented large uniform stimuli of 6 deg visual angle diameter on a gray FSB. For the chromatic condition, we used stimuli that were either green, red, or blue, at three different luminance levels (which are shown in Figure 4). For Figure 1, only the chromatic conditions with the highest available luminance level were used, approximately corresponding to the maximum possible luminance level for the blue primary. For the achromatic condition, we used either black (minimum luminance) or white (maximum luminance) stimuli.

The background was of an intermediate gray value that allowed for good eye tracking quality (see Table S1 for all luminance, RGB and CIE values). Stimulus duration was 3.3 s.

#### Dataset 2

For Figure 2, i.e. the size tuning paradigm, we presented a smaller (either 0.5, 1, or 2 deg) stimulus and a larger (6 deg) surface stimulus in the same trial sequentially, with each stimulus presented for only 0.6 s. In each trial, either the smaller (“small-first”) or largest (“large-first”) surface was presented first. In addition, we used an “edge” condition in which the selected multi-unit’s RF was centered around the vertical edge of the 6 deg stimulus, again followed or preceded by the standard full condition (Figure S4). The colors used were red, blue and green (at the same luminance intensities shown in Figure 1), black and white, and in case of monkey H, also orange, cyan and magenta hues.

#### Dataset 3

For Figure 3, we used only red, green and blue hues (with the same luminances as the maximum luminant red, green and blue used in *Dataset 1,* Figure 1). We presented three stimulus conditions: The uniform surface, the “annulus” and the “blob” condition (Figure 3). Stimuli in annulus or blob conditions were of the same size as the uniform surface, but the center 1 deg of the surface was either surrounded by a thin (0.25 deg) annulus of one of the other, equiluminant, hues, or filled completely with one of the other hues (Figure 3). For each surface of a given hue, there were therefore two “annulus” and “blob” conditions with the two remaining colors (Figure 3). In the analysis, we averaged over all the color combinations for a given condition, and compared the three main conditions. For monkey H, we additionally recorded two sessions with maximally luminant instead of equiluminant hues. Note that this generated strong luminance contrast changes between the colors, but yielded qualitatively similar results. This indicates that the observed effects do not depend on equiluminance, a condition that may occur rarely in nature. Because results were qualitatively similar, we pooled these sessions with the remaining 5 sessions of this animal. We used stimulus presentation times of 1.3-3.3 s. The first 1.3 s were analyzed, as in Figure 1.

#### Dataset 4

For Figure 4A, we recorded “rainbow” sessions in which surfaces (again 6 deg diameter size) of different colors were presented at the maximum possible luminance. We sampled the visible light spectrum linearly in 15 steps of equal size in terms of wavelength, with the MATLAB (MathWorks, Inc.) internal function spectrumRGB.m. Note that the monitor cannot produce line spectra, but can only approximate the corresponding hues through mixing of RGB channels (see e.g. Figure 7A for a yellow hue). We additionally included brown and pink (extra-spectral) hues (see Table S1 and Figure Figure S4A) and achromatic stimuli. For the analyses shown in Figure 4B, we used *Dataset 1*.

#### Dataset 5

For Figures 6 and 7, we used FSBs of various hues. The backgrounds used were red, green, blue and yellow at maximum possible luminance, as well as black, white and gray, presented at the same luminance intensities as in the other datasets. Surface stimuli of 6 (monkey H) or 8 (monkey A) deg diameter in size were used. The size was slightly increased for monkey A to place the edge of the surface stimulus further from the most peripheral RFs. The hues used for the surface were identical to the ones used for the FSBs. In addition, we presented chromatic surfaces with reduced values, namely red, green and blue with the same luminance levels as in Figure 1, and a brown surface. All possible combinations of surface and FSB hues were shown. All other presentation parameters were kept as for *Dataset 1*.

For all stimulus paradigms for monkey A, and in *Dataset 5* for monkey H, there was a post-stimulus period of 0.7 s (0.5 s in monkey H) after the offset of the stimulus, during which the monkeys was required to maintain fixation. For monkey A, the fixation color would change after this period and the monkey had to respond to this change with the release of a lever, whereupon the fixation spot was removed. Presentation of different stimulus conditions was in a pseudo-random order. Typically, 15-20 correctly performed repetitions per condition and session were collected (see also Figure captions for trial numbers).

### Data analysis

#### Preprocessing

Data were analysed in MATLAB using the FieldTrip toolbox (Oostenveld et al., 2011). Only correctly performed trials were analyzed. LFPs were derived from the broadband signal using MATLAB’s decimate.m function, by low-pass filtering with a cutoff frequency of 24414.0625/24/2 Hz (FIR Filter with order 30) and downsampling to 24414.0625/24 Hz. Line noise was removed using two-pass 4 ^th^ order Butterworth bandstop filters between 49.9-50.1, 99.7-100.3 and 149.5-150.5 Hz. LFPs had a unipolar reference scheme described in *Recordings.* Explorative analyses with local bipolar derivations, obtained by subtracting the signals from immediately neighboring electrodes from each other, yielded comparable results (data not shown). MU signals were derived from the broadband signal through bandpass filtering between 300 and 6000 Hz (4 ^th^ order butterworth), rectification, and applying low-pass filtering and downsampling the same way as for the LFPs. For the calculation of rate modulations, this MU signal was smoothed with a Gaussian kernel with an SD of 20 ms. Qualitatively similar results were obtained using thresholded multi-unit data. We used this MU signal for all analyses in the main text, as in previous studies by other labs (Legatt et al., 1980; Schmid et al., 2013; Self et al., 2013; Xing et al., 2012).

#### Receptive field estimation

Receptive fields were mapped with moving bar stimuli (spanning the entire monitor). Moving bars (width 0.1 deg, speed 10/17 deg/s) were presented in 8 orientations for monkey H and 8-16 orientations for monkey A, each for 10-20 repetitions. Mapping sessions were intermittent for monkey H and typically daily for monkey A, to confirm stability of the recordings. MU responses were projected onto the stimulus screen, after shift-correction by the response latency that maximized the back-projected response. MU responses were then fitted by a Gaussian function. This Gaussian was used to extract the 10th percentile and the 90th percentile, and this was done separately for each movement direction. Across the 16 directions, this yielded 32 data points, which were fit with an ellipse. This ellipse was defined as that MU’s RF.

#### Electrode selection

We included all electrodes for analysis that met the following criteria: (1) the MU showed a response to RF stimulation that was at least two SDs above stimulation outside the RF. (2) The MU response during the response period (0.05-0.15 s) of at least one condition of the respective dataset was at least 2 SD above the corresponding baseline (-0.1-0 s). In case of Figures 2-3, it was additionally required that the RF center of the MU was within 0.5 deg of the stimulus center. In the remaining figures, it was required that the RF center was within the surface stimulus.

#### Estimation of LFP power spectra

For Figures 1, 3-4 *and* 6-7, the baseline period was the last 500 ms before stimulus onset, and each stimulation period yielded two non-overlapping epochs of 500 ms (0.3-1.3 s period). For Figure 2, due to the short presentation times, we used epochs of 300 ms (300-600 ms after the onset of the stimulus, and for baseline 300 ms before stimulus onset). LFP epochs were multiplied with discrete prolate spheroidal sequences (multi-tapers for ±5 Hz smoothing), Fourier transformed and squared to obtain LFP power spectral densities (for a recent discussion on spectral estimation see Pesaran et al. (2018)). For Figure 5, we used windows of 0.3 s length, slid over the data in steps of 50 ms. Data were multiplied with a Hann taper before Fourier transformation.

#### Normalization of LFP power spectra

To show LFP power changes, we computed relative power spectra by dividing single-trial power spectra from the stimulation period by the average power spectra across conditions and trials from the baseline. This was shown as a fold-change in all figures showing relative changes except for Figure 5 TFRs, where for visualization purposes, we transformed this into dB units.

To investigate absolute LFP power (without reference to the baseline), we normalized power spectra per electrode by the total power above 25 Hz in the baseline condition. This normalization reduced variance or scaling in the LFP power spectra across sessions and animals before averaging. By normalizing both the baseline and the stimulus period by the same normalization factor, we could still examine changes in raw LFP power across conditions, for each frequency bin separately. This would not have been possible if we had normalized the LFP power spectrum in a given condition by the total power across frequencies in the same condition. These power spectra were averaged across the selected channels (except for single-channel analyses as in Figure S2).

#### Quantification of LFP gamma-band amplitude

Quantification of the differences in gamma-band amplitude between conditions is in general a difficult problem because changes in firing rate can cause broad-band shifts in the LFP power spectrum, and because spikes can “bleed-in” at higher LFP frequencies (Buzsaki et al., 2012; Miller et al., 2009; Pesaran et al., 2018; Ray and Maunsell, 2011). We developed an algorithm to extract gamma-band amplitude in order to address these problems (see Figure S1 for an illustration). We present two versions of this algorithm that are used for separate figures, and are based on constructing a polynomial fit of the LFP spectrum which was detrended in two separate ways. The first algorithm had the following structure:

1. Power spectra were log-transformed and the frequency axis was also sampled in log-spaced units to avoid overfitting of high-frequency datapoints. All subsequent polynomial fits were performed on the 20-140 Hz range.
2. We used the change in stimulus-induced LFP power versus the common baseline (see above), expressed as ΔP =log(*P*_*stim*_) — log(*P*_*base*_)
3. To determine the polynomial order, we used a cross validation procedure to prevent overfitting. A random half of the trials was used for the fitting and deemed the “training set”. The remaining trials were the “test set”. Polynomials of order 1-20 were fit to ΔP as a function of frequency for the “training set”, minimizing the mean squared error. We then computed the mean squared error using the same polynomial fit on the “test set” for each of the 20 orders. This procedure was then repeated for multiple (50) iterations, with a random half of the trials selected for each iteration, and for each iteration, the best-performing order was retained.
4. A polynomial with the median of the best-performing orders was then fit to the complete set of trials.
5. On the polynomial fit, local maxima and minima in the 30-80 Hz range were identified. The peak gamma frequency was the location of the maximum. The band-width of gamma was estimated as twice the distance between the frequency of the maximum (*F*_*max*_) and the frequency of the first local minimum to the left of the maximum (*F*_*min*_), i.e. *b* = *2F*_*max*_ *- F*_*min*_ (Figure S1). The gamma amplitude was then assessed from the difference between the value of the polynomial fit at the maximum and the average of the polynomial fit at *F*_*min*_ and *F*_*max*_ + *F*_*min*_ (Figure S1).
6. This difference was taken in log-space (because the power spectra were originally log-transformed) and then transformed to a fold-change.

If firing rate changes relative to baseline (or between conditions) were very strong, e.g. with small stimuli, this fitting procedure occasionally ran into problems, because relative LFP power spectra showed broad increases that were likely due to non-rhythmic processes like spikes or postsynaptic potentials (see Figure 3 for an example of this effect). In addition, in Figure 6 and 7, because we used background stimuli of different hues, a “neutral baseline” like the gray background screen was not always available. In these cases we modified the second step of this algorithm. Instead of computing the change in LFP power relative to baseline, we performed a 1/F ^n^ correction on the raw LFP power spectrum. The 1/F ^n^ correction was performed by fitting an exponential to the LFP power spectrum, excluding data points in the typical gamma range of 30-80 Hz. Note that we fitted an exponential function because in many cases, bleed-in of spiking energy in the LFP caused a departure from a linearity in the log(power) versus log(frequency) graph (see also Haller et al. (2018); Shirhatti and Ray (2018)). We visually inspected the fits for a large number of spectra and compared this also to a procedure with a mixture of a linear fit and a Gaussian fit to the log(power) versus log(frequency) graph, which had substantially more problems in dealing with spike-bleed at high frequencies, as well as with additional peaks (potentially harmonics) at higher frequencies (e.g. for the red surfaces) (data not shown).

#### Spike-field coherence

For spike-field coherence, we used only electrodes selected by the procedure described above. In addition, for LFP-MUA pairs, we required that the electrodes were direct neighbors in the grid, and in the case of monkey H, given that the microelectrode array had two fixed depths, were of the same depth. Spike-field phase-locking was computed as follows. We estimated the cross-spectral density between LFP and MU signal for each trial separately (cross-spectra) using the same spectral estimation settings as for the LFP power spectrum. This yielded one cross-spectrum per trial. We then normalized the cross-spectrum per trial by its absolute values, to obtain the cross-spectral phases (without amplitude information). We used those normalized cross-spectra to compute the Pairwise Phase Consistency (PPC), using FieldTrip (Oostenveld et al., 2011). This measure has the advantage that the bias by trial count, inherent to e.g. the spectral coherence, is avoided (Vinck et al., 2010b). For a given MU site, the PPC values were then averaged across all the combinations with LFPs from the other selected channels. Note that MU-LFP combinations from the same electrode were excluded to avoid artifactual coherence due to bleed-in of spikes into the LFP (Buzsaki et al., 2012; Ray and Maunsell, 2011). Because of the distance between electrodes (at least 400 micrometer), this was not an issue for MU-LFP combinations from different electrodes.

The standard error of the PPC was estimated across sessions. This was different from SE estimation for power and rate, which used the bootstrap (see below). Bootstrap estimates are problematic for PPC because bootstraps contain repetitions of identical trials, which trivially yield high coherence values.

#### Rate modulation

Rate modulation was computed as log10 *M*_*stim*_*/M*_*base*_, where *M*_*stim*_ and *M*_*base*_ represent the MU firing activity in the stimulus and baseline period, respectively. To quantify surround suppression, we took the differences of these rate modulation indices between small and large stimulus size conditions.

### Modulation index of fold-changes

To quantify the modulation of LFP gamma-amplitude (expressed as fold-change) between conditions (Figure 5, 6), we computed a modulation index as (*A*-*B*)/(*A*+*B*), where *A* and *B* are the gamma-amplitudes in the two conditions, taken as the fold-change minus 1. Note that the fold-change was extracted using the polynomial fitting procedure described above, and a fold-change of 1 indicated the absence of a gamma peak.

### Statistics

Error bars or shaded error regions correspond to ± one standard error of the mean (SEM). SEM was estimated using a bootstrap procedure, with the exception of spike-field coherence (see above). For the b-th bootstrap out of *B* = 1000 bootstraps, *b* = 1,…, *B,* the following was done. For each condition in a given session, with a set of *N* trials 𝒯, we took a random set of *N* trials from *𝒯* with replacement, yielding a new set of trials *𝒮* _*b*_. For that sample of *N* trials *𝒮* _*b*_, we then computed the statistic of interest. For LFP signals, we then computed the average statistic in a given session over all channels, then averaged over sessions, and then monkeys. The rationale behind averaging across all LFP channels was that these signals are likely highly statistically dependent because of volume conduction among the relatively closely spaced electrodes. For MU signals, we computed the average statistic of interest across sessions per MU site separately, and then averaged across all recording sites. The standard error of the mean was then defined as the standard deviation over the *B* average statistics, as is common with bootstrapping procedures.

We used the bootstrap distributions for inference on fold-change estimates or fold-change modulation indices between conditions, as well as differences in peak gamma frequency. In this case, we computed for each bootstrap the difference between average statistics for two conditions, and then tested whether this distribution was different from zero (with Bonfer-onni correction for number of comparisons).

For frequency- or time-resolved differences (in absolute and relative LFP power spectra and rate modulation scores), we used multiple-comparison corrected permutation tests: In this case, we shuffled the trials between two conditions per permutation *P* times, and then constructed a permutation distribution of average absolute differences between conditions. We equalized trial numbers for each comparison, for example between chromatic/achromatic conditions or the different stimulus sizes. We then compared the observed difference between average statistics against this permutation distribution. For multiple-comparison correction, we used the procedure from Korn et al. (2004), which is based on the sorted distribution of absolute differences, with alpha and false discovery rate values of 0.05. In this iterative procedure, values in the observed distribution exceeding the 95th percentile of the *P* maximal values of each permutation distribution (critical value) are deemed significant. Significant values are removed from the observed distribution, and the same positions are removed from all *P* permutation distributions. Values in the observed distribution exceeding the critical value based on these permutation distributions are then iteratively collected until no value in the observed distribution exceeds the critical value. Note that statistical parameters are reported mostly in the figure captions.

### Quantitative model for dependence of gamma-band amplitude on background stimulus

Cone data were extracted from Harosi (1987) (bleaching difference corrected spectra). Polynomials of order 7 were fit to these curves. The cone response curves were then normalized to the maximum. We measured the spectral energy of each color as well as black, white and gray (Ocean Optics WaveGo; XWAVE-STS-VIS-RAD). The spectral energies of the colors were normalized to unit mass. For gray, we added the normalized energies of R, G and B and multiplied with the energy ratio of gray over white. We then convolved the cone response curves with the normalized spectral energies to determine how strongly each background adapts the three cones. Regression models were then fit as explained in the Results text and caption of Figure 7. SEM for regression coefficients are obtained by the same bootstrap procedure as described above.

## Supplementary Material for: Surface color and predictability determine contextual modulation of V1 firing and gamma oscillations

**Figure 1:**
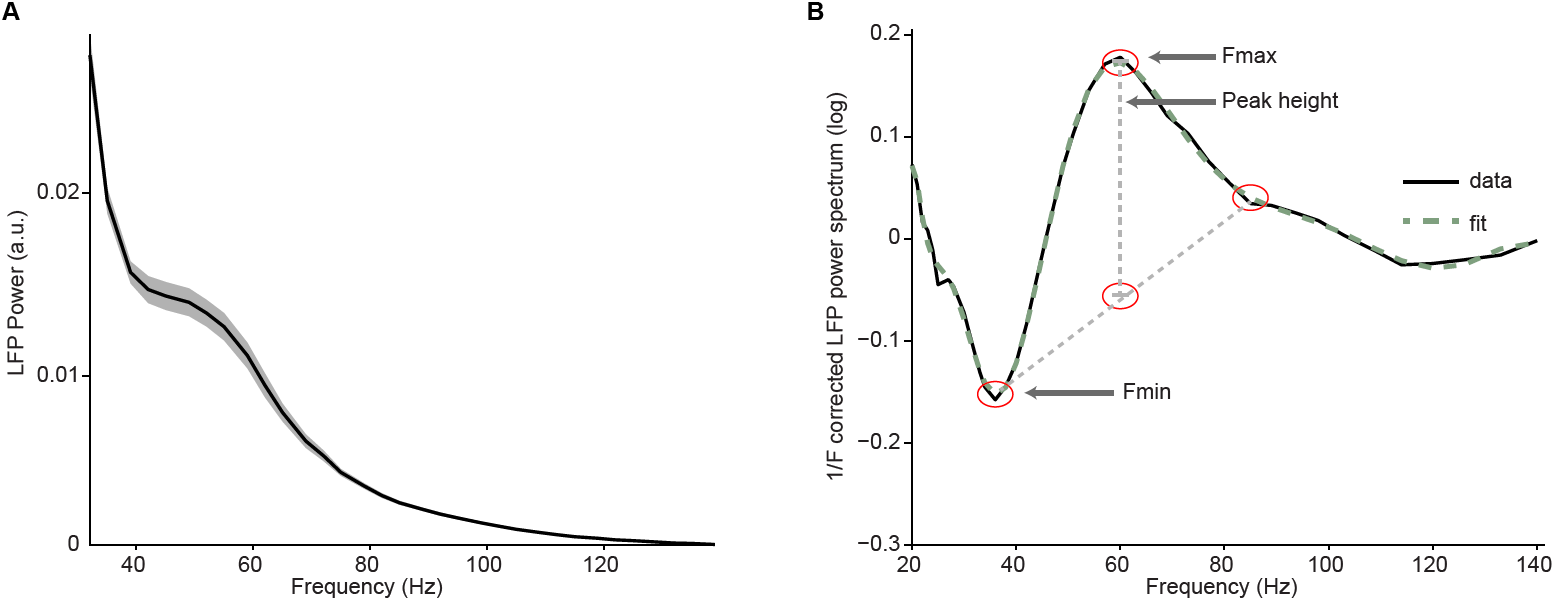
Illustration of fitting procedure. (A) Average LFP power spectra for a large chromatic condition of an example session used in Figure 2. LFP spectra for all conditions were normalized to the summed power (>25 Hz) for the baseline (gray) condition (see Methods). (B) Log-transformed, 1/F ^n^ corrected spectra (solid line) and their fit (dashed line). Peak height was determined as the difference between the peak value at location Fmax and a baseline estimate based on the average of the power at location F_min_ and F_min_+2*(F_max_-F_min_), the estimate of peak width.

**Figure 2:**
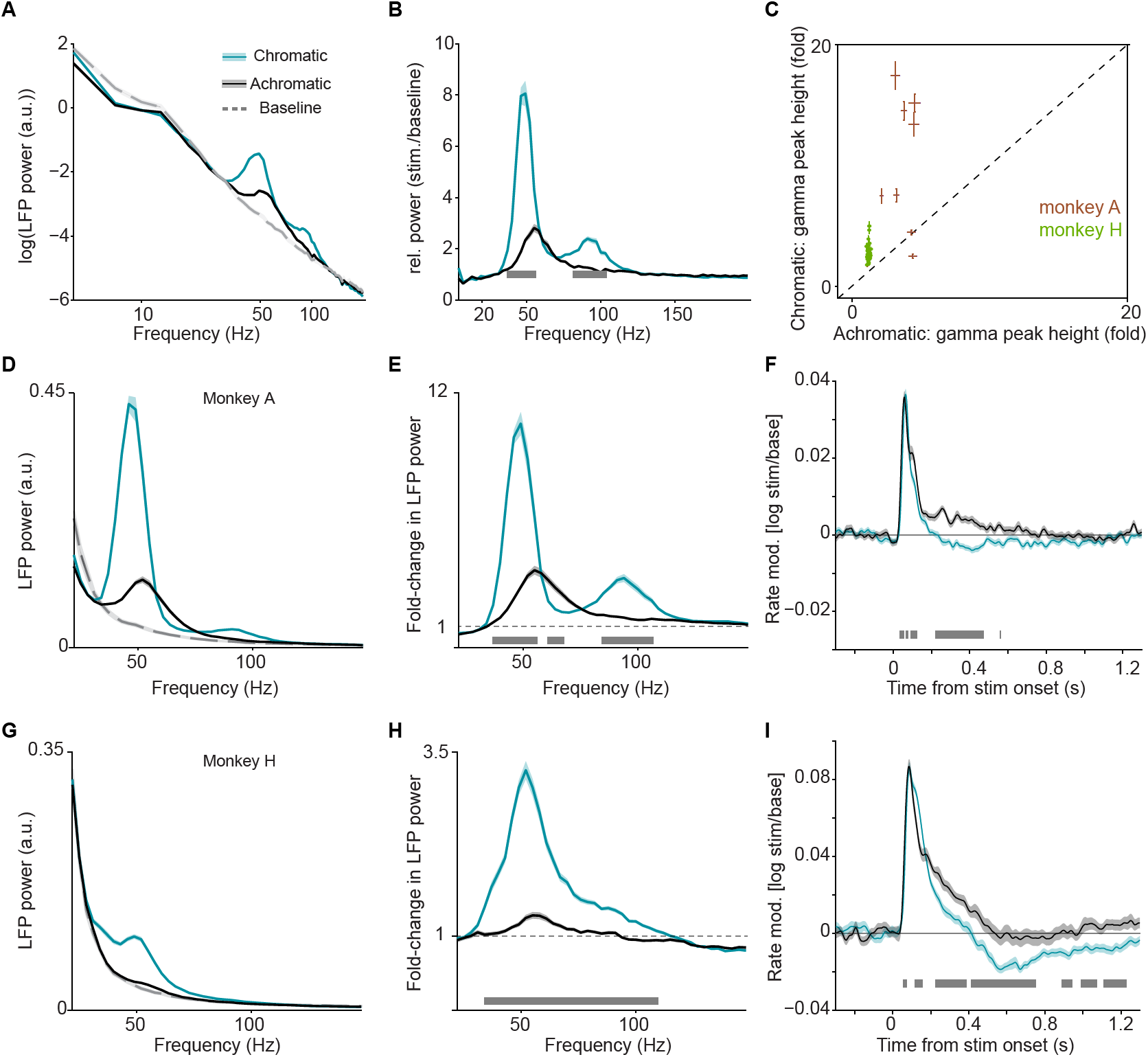
Analysis of LFP and multi-unit activity in response to large, uniform surfaces, related to Figure 1 in the main text. (A) Average LFP power spectra for chromatic (turquoise), achromatic (black) and baseline (gray) conditions. LFP power was estimated using Discrete Fourier Transform of non-overlapping snippets of 500 ms with a Hanning taper. LFP spectra for all three conditions were normalized to the summed power (>20 Hz) for the baseline (gray) condition (see Methods). (B) Average change in LFP power, expressed as fold-change, relative to baseline. (C) Scatter-plot for all the LFP recordings sites in two monkeys, showing the gamma-band amplitude (expressed as fold change) in chromatic and achromatic conditions. (D) Average LFP power spectra for chromatic (red), achromatic (black) and baseline (gray) conditions for monkey A, using same estimation settings and normalization for power spectral density as in Figure 1 of main text. (E) Average change in LFP power, expressed as fold-change, relative to baseline. (F) Modulation of firing rate relative to baseline, expressed as log_10_ (stim/base), for monkey A. (G-I) as (D-F), but now for monkey H. (A-I) Shadings and error bars indicate standard errors of the means (see Methods). Gray bars at bottom of figure indicate significance bars, obtained from permutation testing with multiple comparison correction across all frequencies and time points.

**Figure 3:**
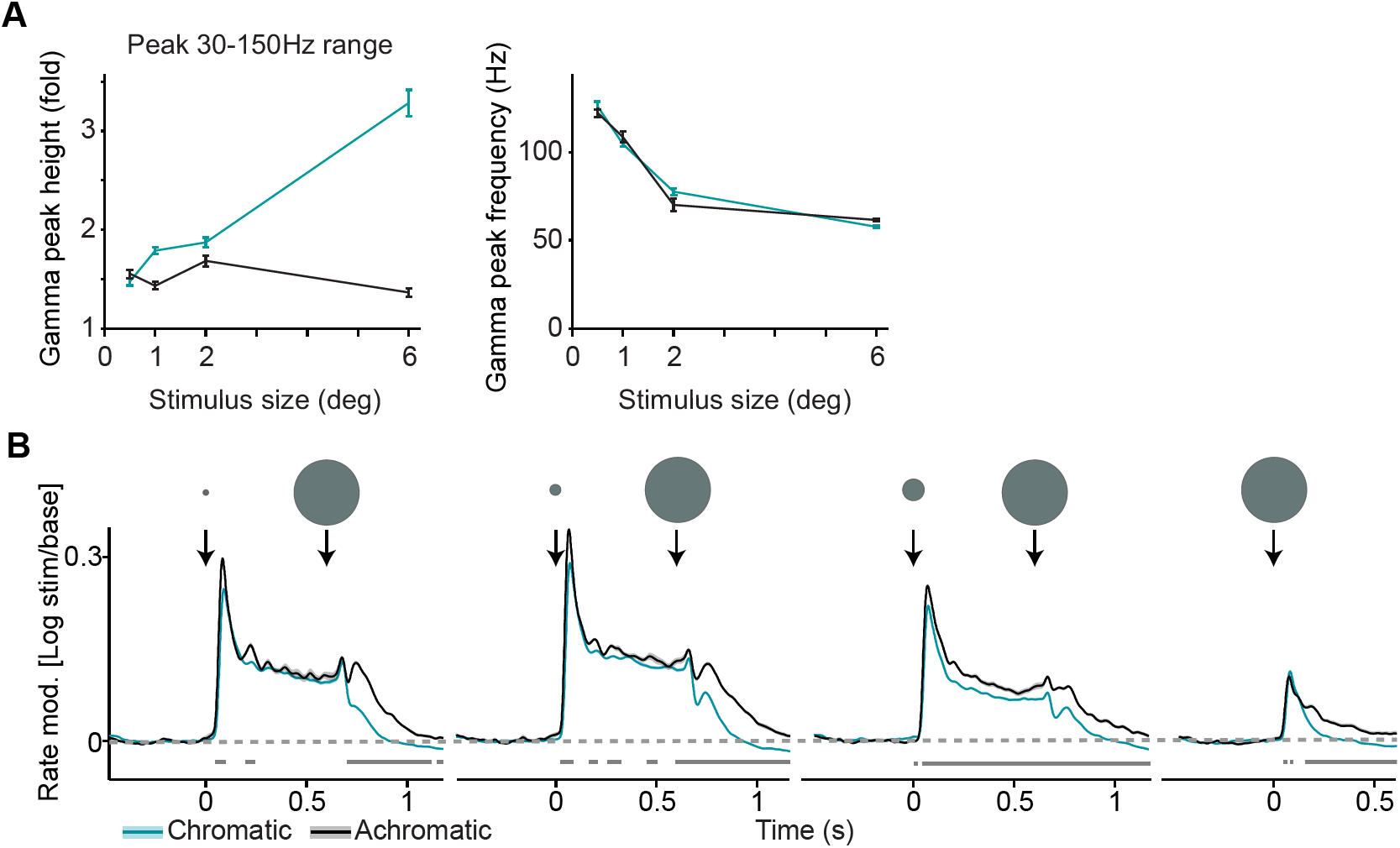
Analysis of LFP and multi-unit activity in response to stimuli of varying size. (A) Gamma-band peak amplitude and peak-frequency as a function of size, estimated using a polynomial fitting procedure between 30-150 Hz. A wider range instead of the standard 30-80 Hz range was used here, to also capture peaks >100 Hz, which is far outside the typical range of classical visual gamma range. This activity may reflect spike bleed-through, which is beyond the scope of the present study. (B) Each trial contained a sequence of two stimuli, either the small stimulus first, or the large stimulus first (see Methods). Here we show the first type of sequence to illustrate the onset of a surround when the stimulus covering the classical RF is already present. Modulation of firing rate relative to baseline, expressed as log_10_ (stim/base), for different sizes and chromatic/achromatic conditions.

**Figure 4:**
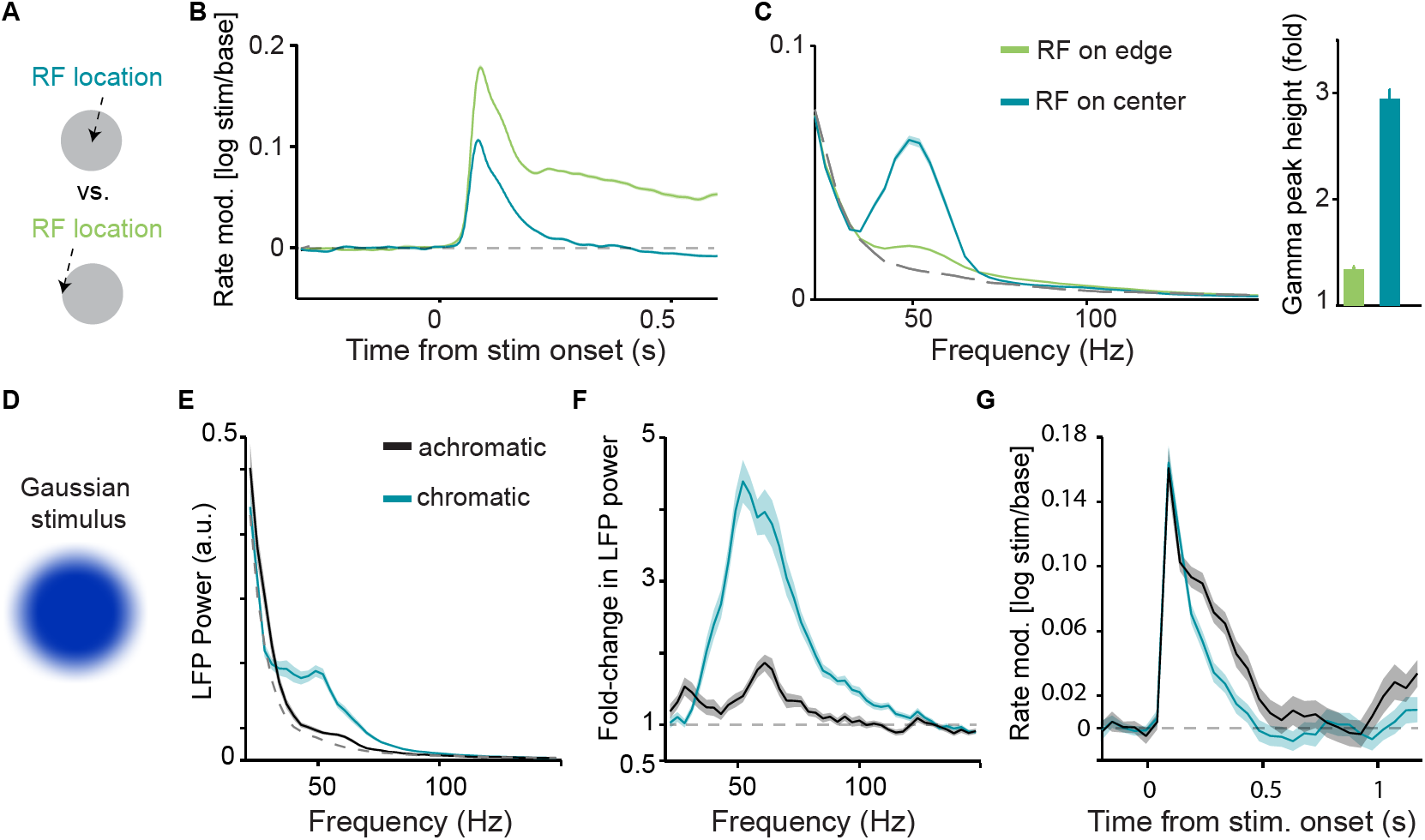
Comparison of V1 responses between RF centered on stimulus center and edge, and V1 responses to Gaussian stimuli. (A) Illustration of paradigm (*Dataset 2*). Uniform surfaces were either centered on the unit’s RF, or the edge of the surface was centered on the unit’s RF. (B) Modulation of firing rate relative to baseline, expressed as log_10_ (stim/base). (C) Average LFP power spectra, using the same analysis time window and spectral estimation parameters as in Figure 2 of main text, comparing “RF-on-center” and “RF-on-edge” conditions. Dashed gray line corresponds to baseline (gray background screen). Right: Gamma-band amplitude (expressed as fold-change) for the two conditions. Gamma-band amplitude was significantly higher for “RF-on-center” condition (P<0.05, bootstrap test). (D)-(F): Single session (from monkey H) illustrating responses to Gaussian surface stimuli. Stimuli were otherwise the same as *Dataset 1.* Gamma oscillations were not abolished by removal of the sharp stimulus edge. (D) Example Gaussian stimulus that had a blurred edge. (E) Average LFP power spectra for chromatic (turquoise), achromatic (black) and baseline (gray) conditions, computed as in Figure S1. (F) Average change in LFP power, expressed as fold-change, relative to baseline. (G) Modulation of firing rate relative to baseline, expressed as log_10_ (stim/base), for monkey A.

**Figure 5:**
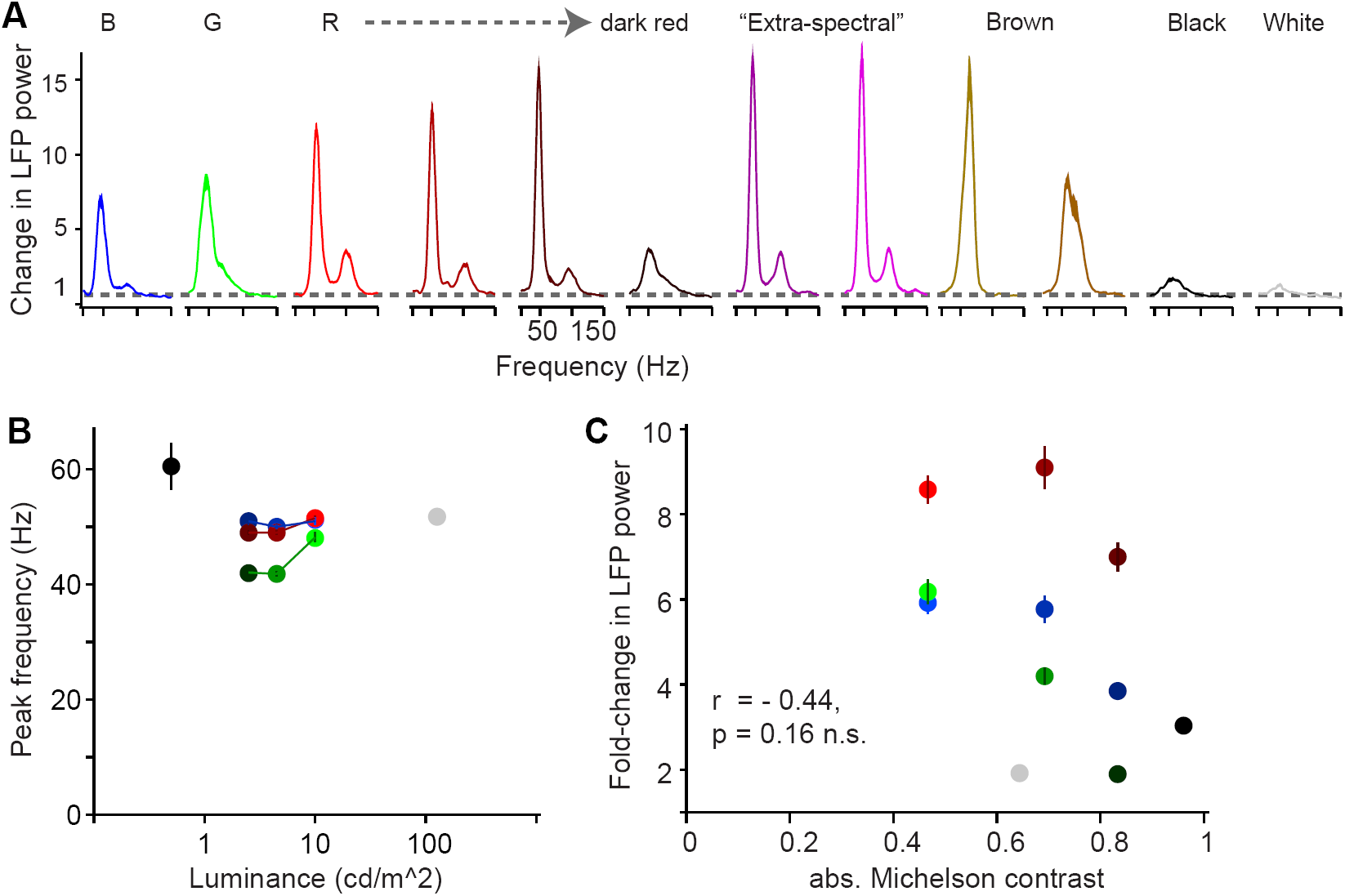
Dependence of gamma LFP power on the surface hue: further hues, peak frequency dependence and control for luminance contrast. (A) LFP power spectra as in Figure 4A, for additional hues. The pure blue, green and red stimuli are shown for reference, followed by red surfaces of decreasing luminance and the brown surface stimuli. See Table S1 for CIE and luminance values. (B) Peak frequency estimates (Hz) based on cross-validated fitting procedure (see Methods) for the surface stimuli of Figure 4B. (C) A regression of absolute luminance contrast (Michelson contrast) against gamma peak height (fold-change) showed no significant relationship (p=0.23). Note that since there was relatively little gamma power for achromatic, high-contrast stimuli, if anything there would be a negative relationship between luminance contrast and gamma power, the very opposite of findings about gamma power for achromatic gratings. Also note that for red stimuli, gamma power appears to follow a U-shape with decreasing luminance and increasing contrast (see also FigS4A).

**Figure 6:**
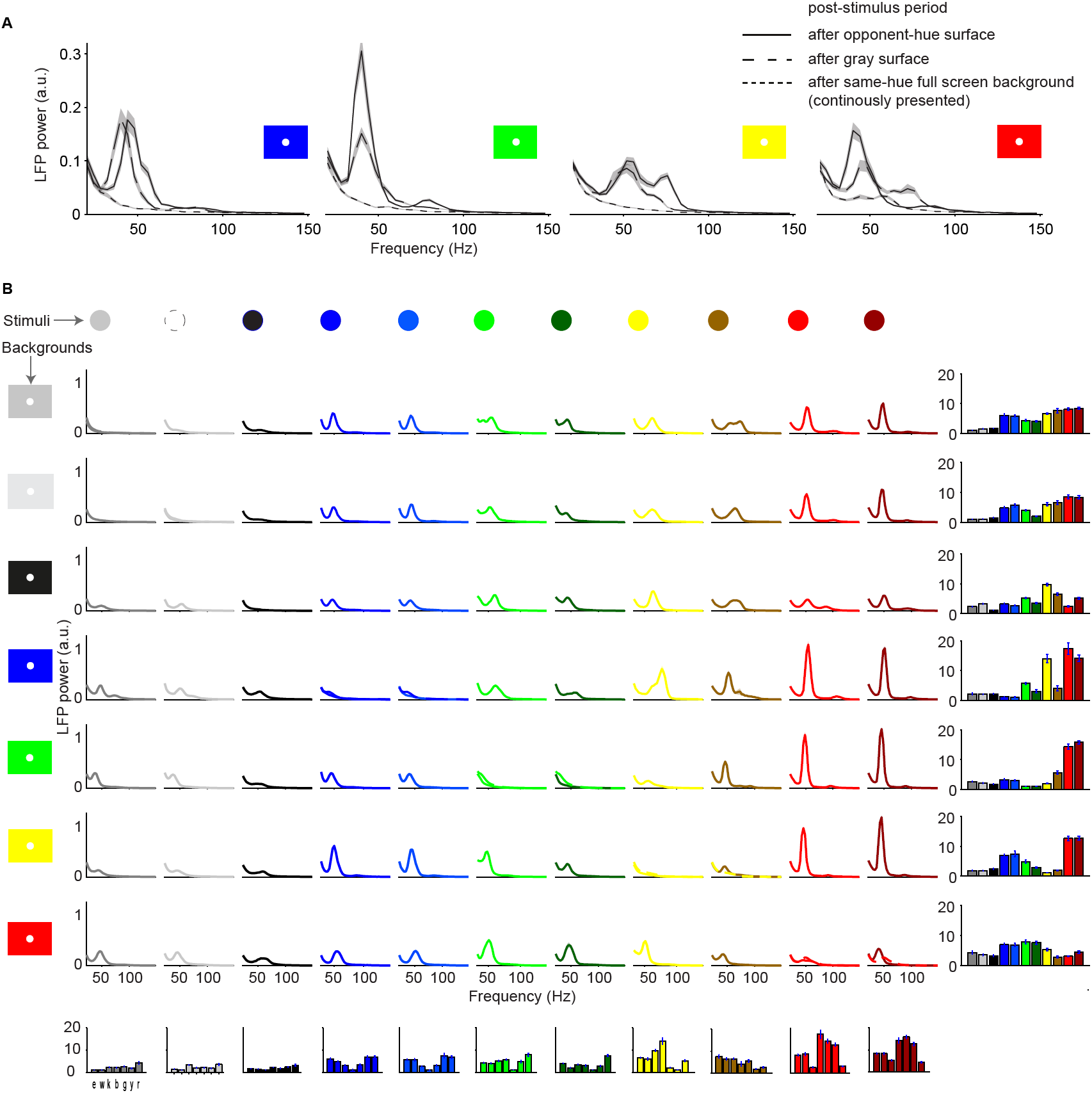
Dependence of gamma LFP power on the combination surface hue and background stimulus. (A) LFP power spectra for the post-stimulus period for the four chromatic background hues (3.5-3.8 s, excluding the initial transient response after stimulus offset at 3.3 s). Clear gamma-band responses for full-screen surfaces after both gray and opponent-hue surface presentation in the stimulus period. (B) We show here all the condition combinations for *Dataset 5.* Different rows correspond to different stimulus background conditions. The color of the background is indicated by the background stimulus shown on the left. The second row corresponds to a white background. Different columns correspond to different stimulus hue conditions, which are indicated by the color of the lines in each graph. Each graph depicts the average LFP power spectrum, using the same estimation parameters as in Figure 6 of the main text. Bar graphs on the bottom show the gamma peak amplitude (fold-change) for the different backgrounds, separate for each surface hue. Bar graphs on the right show the gamma peak amplitude (fold-change) for the different surface hues, separate for each background condition.

**Table 1:** RGB values, luminances (cd/m ^2^) and CIE values used in this study. Luminances and CIE values were measured with a Konica Minolta CS-100A chromameter, CIE values refer to the 1931 2 degree standard observer. Standard black, white and gray used across datasets are in rows 1-3.

| Dataset | R | G | B | Lum | CIE<sub>x</sub> | CIE<sub>y</sub> |
| --- | --- | --- | --- | --- | --- | --- |
| 1 | 0 | 0 | 0 | 0.55 | 315 | 358 |
| 1 | 128 | 128 | 128 | 27.5 | 347 | 357 |
| 1 | 255 | 255 | 255 | 127 | 343 | 372 |
| 1 | 149 | 0 | 0 | 9.69 | 624 | 340 |
| 1 | 0 | 99 | 0 | 10 | 272 | 555 |
| 1 | 0 | 70 | 255 | 10.7 | 161 | 125 |
| 1 | 104 | 0 | 0 | 4.77 | 608 | 311 |
| 1 | 0 | 69 | 0 | 4.41 | 273 | 532 |
| 1 | 0 | 49 | 179 | 4.97 | 163 | 121 |
| 1 | 75 | 0 | 0 | 2.54 | 574 | 341 |
| 1 | 0 | 50 | 0 | 2.83 | 287 | 513 |
| 1 | 0 | 35 | 128 | 2.54 | 169 | 129 |
| 4 | 32 | 0 | 98 | 1.35 | 207 | 129 |
| 4 | 104 | 0 | 255 | 13.4 | 206 | 119 |
| 4 | 96 | 0 | 255 | 12.6 | 198 | 114 |
| 4 | 0 | 0 | 255 | 9.42 | 153 | 91 |
| 4 | 0 | 81 | 255 | 14.8 | 162 | 135 |
| 4 | 0 | 198 | 218 | 49.4 | 218 | 356 |
| 4 | 0 | 255 | 72 | 85.1 | 277 | 573 |
| 4 | 0 | 255 | 0 | 86.2 | 281 | 589 |
| 4 | 66 | 255 | 0 | 89.8 | 293 | 581 |
| 4 | 255 | 247 | 0 | 110 | 430 | 486 |
| 4 | 255 | 154 | 0 | 58.5 | 520 | 417 |
| 4 | 255 | 0 | 0 | 32.3 | 641 | 343 |
| 4 | 179 | 0 | 0 | 14.1 | 631 | 341 |
| 4 | 94 | 0 | 0 | 4.01 | 593 | 338 |
| 4 | 43 | 0 | 0 | 1.12 | 494 | 349 |
| 4 | 159 | 127 | 0 | 28 | 457 | 453 |
| 4 | 159 | 95 | 0 | 20.1 | 510 | 417 |
| 4 | 159 | 0 | 159 | 13.3 | 411 | 218 |
| 4 | 223 | 0 | 223 | 28.2 | 406 | 215 |
| 5 | 255 | 0 | 0 | 32.3 | 641 | 343 |
| 5 | 0 | 255 | 0 | 86.2 | 281 | 590 |
| 5 | 0 | 0 | 255 | 9.42 | 153 | 92 |
| 5 | 255 | 255 | 0 | 115 | 425 | 493 |

